# A three-dimensional, discrete-continuum model of blood pressure in microvascular networks

**DOI:** 10.1101/2022.11.23.517681

**Authors:** Paul W. Sweeney, Claire Walsh, Simon Walker-Samuel, Rebecca J. Shipley

## Abstract

We present a 3D discrete-continuum model to simulate blood pressure in large microvascular tissues in the absence of known capillary network architecture. Our hybrid approach combines a 1D Poiseuille flow description for large, discrete arteriolar and venular networks coupled to a continuum-based Darcy model, point sources of flux, for transport in the capillary bed. We evaluate our hybrid approach using a vascular network imaged from the mouse brain medulla/pons using multi-fluorescence high-resolution episcopic microscopy (MF-HREM). We use the fully-resolved vascular network to predict the hydraulic conductivity of the capillary network and generate a fully-discrete pressure solution to benchmark against. Our results demonstrate that the discrete-continuum methodology is a computationally feasible and effective tool for predicting blood pressure in real-world microvascular tissues when capillary microvessels are poorly defined.

## Introduction

The microcirculation is a hierarchical structure of blood vessels consisting of arterioles, capillaries and venules. These vessels are typically classified as those with diameters < 300 *µ*m and by several distinctive characteristics, such as mural cell coverage ^1^ and resistance to flow ^2^. Structurally, arterioles and venules form branching structures which supply and drain an interconnected, mesh-like network of capillaries with diameters < 10 *µ*m. In particular, the effective diffusion distance for oxygen within tissue is limited to approximately 20 - 100 *µ*m, which is influenced by the balance between oxygen transport and consumption ^3^. Therefore, understanding the impact of network architecture on fluid and mass transport in microvascular tissues is essential for a comprehensive understanding of microcirculatory function.

Advances in biomedical imaging of vascular tissues ^4–7^ have paved the way for computational studies which integrate complete vascular architectures with biophysical models to probe the microenvironment *in silico* in a manner that is currently inaccessible in a traditional experimental setting ^8^. Due to the computational challenges of simulating network-scale blood rheology and dynamics using mesh-based methods, many studies apply one-dimensional (1D) Poiseuille flow models. In these models, the vascular network is represented as a graph (see Figure 1A). This approach provides insight into a wide range of biological applications such as cerebral blood flow ^9–12^, angiogenesis ^13,14^, and cancer ^4,15–17^. Nonetheless, with the emergence of whole-organ vascular imaging ^6^, the computational demands of fully-discrete 1D fluid and mass transport models are increasing due to the sheer number of blood vessels in imaged samples (> *O*(10^9^)). Moreover, difficulties such as boundary condition assignment arise when considering incomplete vascular networks which are acquired using a modality with a resolution lower than the size of some of the vessels. For example, photoacoustic imaging is an emerging modality which can image vascular structure and function *in vivo* ^18^ typically at a resolution of 30 − 40 *µ*m, and so is unable to monitor vessels or spatially-resolved flows in individual vessels below this threshold ^19^. This leads to discontinuities in vascular network architecture which increases the number of boundary conditions that require accurate parameterisation. With these considerations in mind, versatile mathematical models are required which are computationally tractable and can bridge the information gap for imaging modalities that cannot resolve vessels or functional parameters down to the micron-scale.

**Figure 1:**
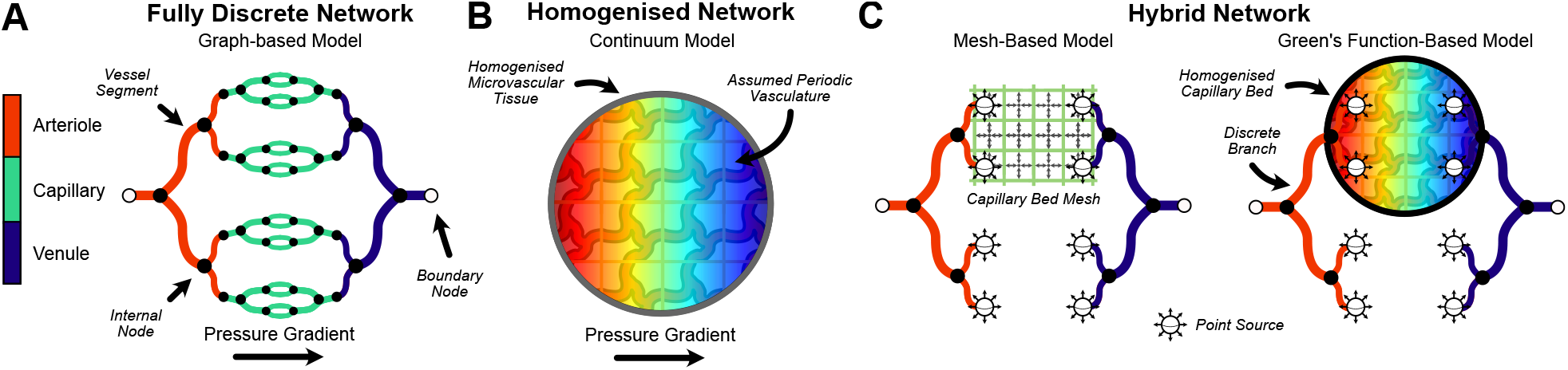
Methods for modelling blood flow in microvascular networks. In healthy tissue, microvasculature can be classified in terms of (red) arterioles, (green) capillaries and (blue) venules. (A) Fully discrete methods can be applied when structural data of the entire blood vessel network is acquired experimental. Here, the vascular network is represented as a graph where nodal pressures, and consequently segment flows, are solved for using a 1D Poiseuille flow model. (B) Mathematical homogenisation can be used when discrete vessel data is unavailable for the entire domain. Darcy’s Law is used to predict tissue-scale blood flow and tissue hydraulic conductivity can be approximated using mathematical averaging techniques, such as those which assume periodic vascular structures. (C) Hybrid models may improve prediction of blood flow heterogeneity across the tissue where structural information of larger vessels is obtained. Here, arteriolar and venular structures are modelled in a discrete fashion, coupled to a (left) mesh or (right) our analytic Green’s function method to solve for blood flow in a capillary continuum.

Mathematical homogenisation provides the opportunity to circumvent the problem of solving tissue-scale fluid and mass transport in the absence of known vascular structure (see Figure 1B). Vascular tissues are represented by a single or coupled continua and modelled as porous media via Darcy’s equation ^20–27^. For example, by assuming two well-separated capillary and tissue length scales, these models enable averaged predictions of pressure and velocity fields with reduced computational cost. However, several challenges remain in their application ^28^. For instance, the range of hierarchical vascular structures may not be fully characterised by two length scales in isolation. Further, these models assume that networks are highly interconnected and so small-scale vessel pressures are correlated which may not be the case, depending on vascular topology. Thus, model parameters, such as the hydraulic conductivity in Darcy’s equation, are difficult to estimate.

Numerous hybrid blood flow models have been formulated which embed larger individual (or discrete) vessels into a homogenised representation of the surrounding vascular tissue ^28–37^. These studies apply a 1D flow model to discrete vessels, coupled to a porous medium Darcy model (see Figure 1C). As a consequence of explicitly representing the hierarchical structure of larger branching vessels and therefore providing a better prediction of flow distribution in these vessels compared to an averaged description, hybrid models may provide a better prediction of blood flow heterogeneity in the homogenised domain. Typically, hybrid models solve for blood flow numerically using semi-analytic or fully finite mesh schemes ^28,30,33,34,36,37^. While these methods have been successful in handling multi-scale problems, they can be computationally intensive, especially for large domains. To mitigate the need for highly resolved meshes, alternative approaches have been proposed that utilise analytical approximations or Green’s functions which couple discrete vessel structures with the Darcy continuum domain (representing the capillary bed) via point sources of flux ^29,35^. In our previous work, this we developed a hybrid, Green’s function model for two-dimensional (2D) networks. Here, the arteriolar network was represented by a vascular graph and coupled to a homogenised capillary bed. Coupling occurred via point sources of flux which were distributed over discs and the venular network was depicted using a single, spatially-uniform network sink term ^35^. Additionally, we ensured conservation of flow between the discrete arteriolar vessel and the venular sink term.

In this study, we develop a 3D hybrid discrete-continuum model for blood flow in microvascular networks. Blood pressure is modelled discretely (1D Poiseuille flow) in the arteriolar and venular branching networks and coupled to a single-phase Darcy description of blood flow in the capillary bed via spherical point sources of flux. We calculate the 3D capillary bed hydraulic conductivity tensor by local averaging of 3D synthetic, periodic micro-cell representations of the capillary vasculature, following established homogenisation approaches. We provide a Green’s function-based solution for the averaged capillary blood pressure in 3D. Compared to similar 2D approaches ^29,35^, our method introduces a capillary network exchange term to address field-of-view limitations in microvascular imaging. Our model does not restrict mass conservation solely to terminal arteriolar / venular branches, but ensures continuous blood flow representation throughout the entire network, even when imaging covers only a sub-volume of the real-world network.

Our model is intended to be used on various microvascular tissues where the structure of the branching arteriolar and venular vessels is known, but the discrete capillary network structure is not. It is important to note that despite multiscale derivations assuming microscale periodicity, Darcy’s Law has been used to predict fluid mechanics at the macroscale for non-periodic structures ^38,39^. Therefore, our model could feasibly be applied to cases where the imaging spatial resolution is above the typical maximum threshold of capillary thickness (10 *µ*m) such that the full arteriolar / venular branching structures are not obtained. In this study, we utilise a fully-resolved vascular network (1.14×1.14×1.72 *µ*m^3^ resolution) from the mouse cerebral medulla/pons to develop our methodology. By classifying the blood vessels of the discrete network into arterioles, capillaries and venules, we use the 3D structural information to generate synthetic micro-cells which, via local averaging, we use to estimate a 3D hydraulic conductivity tensor of the capillary bed. Next, we apply a 1D Poiseuille flow model to the fully-discrete medulla/pons network to generate a blood pressure solution. Using the simulated predictions of capillary hydraulic conductivity and capillary boundary flux (via the discrete flow solution) our hybrid, discrete-continuum model estimates vascular blood pressure across the capillary continuum. The resulting pressure solution is then benchmarked against the fully-discrete medulla prediction and the relative merits of each model’s solution discussed. Finally, we perform sensitivity analysis of key parameters in our hybrid model to explore the role of pivotal structural (e.g., branching order) and functional (e.g., tissue hydraulic conductivity and far-field capillary pressure) parameters in influencing the discrete-continuum model predictions.

## Methods

### Animal model

All animal studies were licensed under the UK Home Office regulations and the Guidance for the Operation of Animals (Scientific Procedures) Act 1986 (Home Office, London, United Kingdom) and United Kingdom Coordinating Committee on Cancer Research Guidelines for the Welfare and Use of Animals in Cancer Research.

### Multi-fluorescence high-resolution episcopic microscopy of Brain

Mouse brains were prepared as described in Walsh et al. ^5^ . Briefly, animals were administered with 200 *µ*l Lectin (Tomato) bound to DyeLyte-649 (Vector UK) (1 mg/ml), administered via tail vein injection and allowed to circulate for 10 min before euthanasia. Perfusion fixation was varried out, the brain removed, and prepared through a graded dehydration and resin embedding process ^5^.

Multi-fluorescence high-resolution episcopic microscopy (MF-HREM) images were collected with voxel size 0.57×0.57×1.72 *µ*m^3^ allowing the entirety of the vascular network to be captured. Images were deconvolved using a diffraction kernel as specified in Walsh et al. ^6^ . The deconvolved images were downsampled to voxel sizes of 1.14×1.14×1.27 *µ*m^3^ and segmented semi-manually in Amira v2020.2, using a magic wand tool ^5^. The segmentation was skeletonised using the Amira autoskeletonise module (with distance-thinning mode enabled) to provide a graph description of the vascular network. Vessels are represented by segments with a defined diameter and length, interconnected by nodes with defined 3D coordinates. A 3D volume of the skeleton was taken from the medulla/pons (located at the base of the brainstem - here on referred as the medulla) and used as the geometrical input into our mathematical model.

In the medulla, larger arteriolar and venular blood vessels penetrate through the tissue with intermediate vessels branching off ^40^. A mesh of capillaries form from the intermediate vessels, connecting the arteriolar and venular branches. We classified vessels according to their type (arteriolar, venular, capillary - see Figure 2) enabling the capillary network to be removed, mimicking an imaging modality constrained by spatial resolution. This left the larger, branching arteriolar and venular networks as discretely-modelled structures in our hybrid approach.

**Figure 2:**
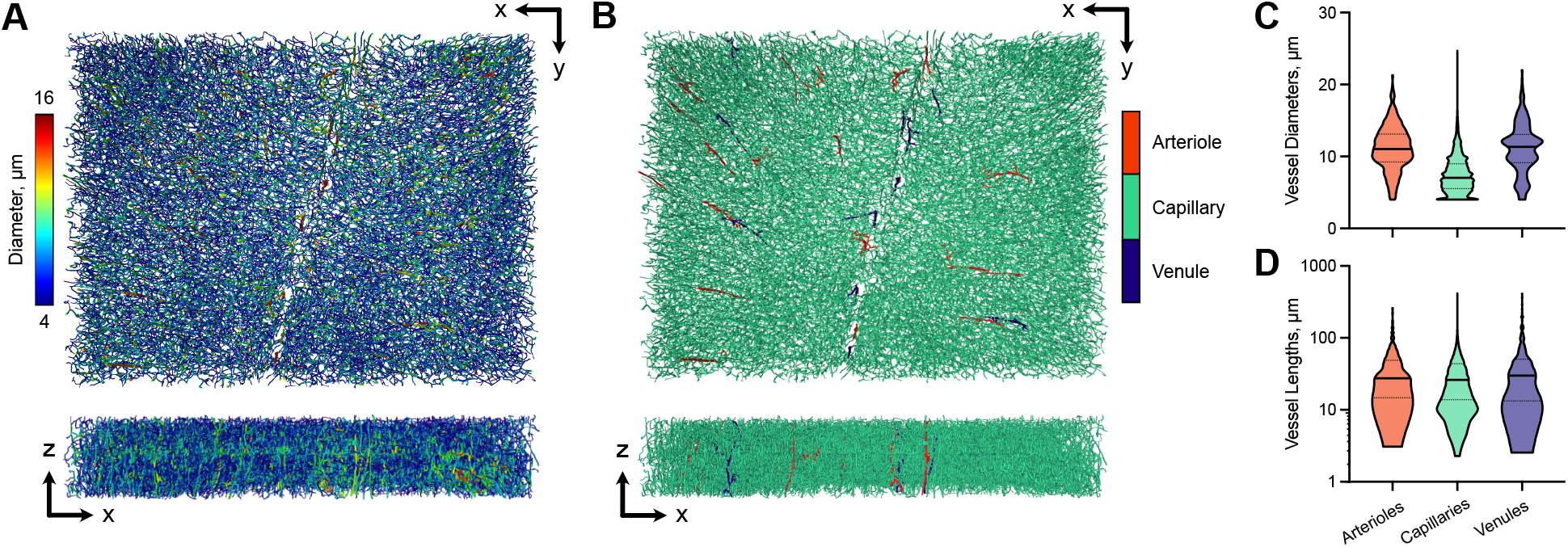
Structure of the vasculature in the mouse medulla. 3D visualisations of the medulla vasculature indicating (A) blood vessel diameters and (B) manually identified arteriolar and venular vessels. (C) Distributions of blood vessel diameters and lengths for arterioles, capillaries and venules in our dataset. A full description of parameters are provided in Table 1.

Vessel classifications are not known *a priori* for the medulla dataset. As such, we pragmatically classified vessels based on vascular morphology ^40^. Penetrating vessels were identified via visual inspection and intermediate branching vessels were located using a breadth-first search with a diameter threshold of 9 *µ*m and all remaining vessels were labelled as capillaries (see Figure 2). We found that setting a 10 *µ*m threshold to identify larger vessels led to inaccuracies due to discontinuities in penetrating arteriolar and venular vessels, likely caused by tissue shrinkage during sample dehydration for MF-HREM imaging. To maintain physiologically-realistic structure without artificially adjusting sizes, a 9 *µ*m threshold was chosen, successfully preserving the structure of penetrating vessels.

Sensitivity of model predictions to the diameter threshold is explored in our results and a summary of the architectural information for the skeletonised vascular network, as well as vessel classifications, is provided in Table 1.

**Table 1:** A summary of vascular network properties for the medulla network. Note, vascular density is defined as the ratio of network volume to tissue volume.

| Metric | Value |
| --- | --- |
| Tissue Dimensions (mm<sup>3</sup>) | $3.09 \times 2.33 \times 0.54$ |
| Vascular Density (%) | 3.10 |
| No. of penetrating vessels | 32 |
| No. of branching vessels | 90,176 |
| No. of branch points (vertices) | 73,175 |
| No. of segments | 319,534 |
| No. of nodes | 302,554 |
| No. of boundary nodes | 21,983 |
| Mean vessel diameter ($\mu\text{m}$) | |
| Arterioles | $11.11 \pm 3.08$ |
| Capillaries | $7.44 \pm 2.49$ |
| Venules | $11.08 \pm 3.43$ |
| Mean vessel length ($\mu\text{m}$) | |
| Arterioles | $37.91 \pm 36.31$ |
| Capillaries | $32.19 \pm 24.96$ |
| Venules | $43.96 \pm 52.20$ |

Simulating blood pressure using a 1D Poiseuille model requires boundary conditions at all the network boundaries. However, functional information for all boundaries, especially for high-resolution data, is rare. As a result, we initially introduce a 1D Poiseuille flow model for incomplete boundary conditions ^41^ which has been applied across a range of real-world tissues ^4,11,17,42^ and estimates unknown boundary data by matching to target mean pressure and shear stress across the network. We apply this model to predict blood pressure across the fully-discrete medulla network to generate our reference pressure solution, as well as predict the flow distribution in branching vessels in our hybrid model.

Next, we present our hybrid model which predicts blood pressure in the homogenised capillary bed coupled to the discrete arteriolar and venular branches. To predict the resistance to flow through the capillary bed, we detail a 3D micro-cell problem to estimate hydraulic conductivity in synthetic networks ^43,44^. This is followed by the computational implementation of the hybrid discrete-continuum model to predict the pressure distribution in microvascular tissue, in the absence of a spatially-resolved capillary network. Finally, we detail our methodology behind boundary condition and parameter assignment for our hybrid, discrete-continuum approach.

### Discrete blood flow model

Vascular networks can be represented by a graph(s) consisting of an interconnected set of nodes, *N*, and segments, *S*. Establishing a consistent flow direction from the start node to the end node of each vessel segment and ensuring flow conservation at blood vessel junctions, the relationship between nodal pressures, *p*_*k*_, and the boundary fluxes, *Q*_0*i*_, is given by

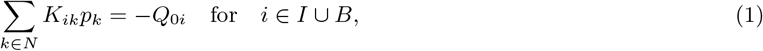

where *I* is the set of all interior nodes and *B* the set of boundary nodes with known flow conditions. If *i* is a boundary node with a known boundary condition, *Q*_0*i*_ is equal to a prescribed pressure or flow value (positive or negative dependent on inflow or outflow, respectively) at node *i*. Note, if a pressure condition is assigned, row *i* of *K*_*ik*_ is set to zero except for start/end nodes of the associated vessel segment, which are set to one. Network conductance to flow, *K*_*ik*_, is defined as

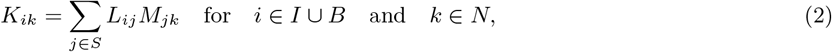

where

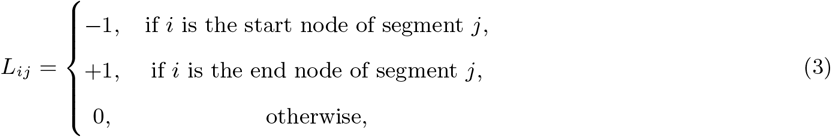

and

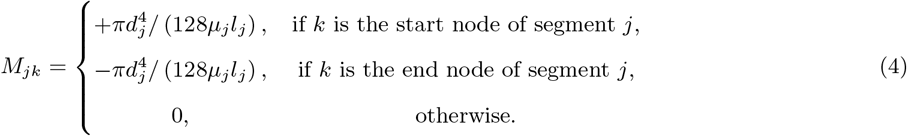

The variables *l*_*j*_, *d*_*j*_ and *µ*_*j*_ denote the length, diameter and effective blood viscosity of segment *j*, respectively. In this study, we define blood viscosity, *µ*(*d, H*_*d*_), as a function of vessel diameter and haematocrit, *H*_*d*_, employing a commonly used empirically-derived blood viscosity law ^45^. For simplicity, we assign a constant haematocrit value of 0.45, however, red blood cell phase separation can be incorporated ^46^ in future work.

When a set of boundary conditions are unknown, we formulate a constrained problem in terms of a Lagrangian objective function ^41^

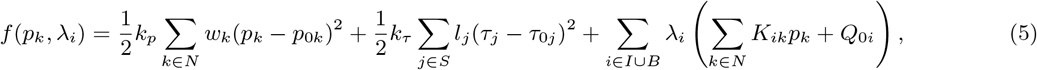

which seeks to minimise the deviation between target nodal pressure, *p*_0*k*_ and segment wall shear stresses, *τ*_0*j*_ subject to the assignment of weighting factors *k*_*p*_ and *k*_*τ*_ . Here, *τ*_*j*_ is the vessel wall shear stress for vessel *j, λ*_*i*_ is a Lagrange multiplier and *w*_*k*_ is a weighting factor and equal to the sum of segment lengths connected to node *k*.

Equation (1) and *∂f/∂p*_*k*_ = 0 can be combined into the following linear system:

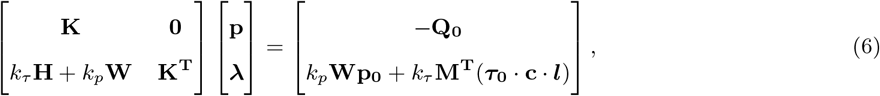

where **K, H, W** and **M** denote the matrix forms of *K*_*ik*_, *H*_*ik*_, *w*_*k*_ and *M*_*ij*_, respectively, noting **W** is a diagonal matrix with entries *w*_*k*_; **p, *λ*, Q**_**0**_, **p**_**0**_, ***τ***_**0**_, **c** and ***l*** are the vector forms of *p*_*k*_, *λ*_*k*_, *Q*_0*i*_, *p*_0*k*_, *τ*_0*j*_, *c*_*j*_ and *l*_*j*_, respectively. Here, *H*_*ik*_ is defined as

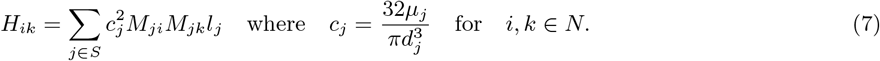

The linear system (6) can be solved for unknowns *p*_*i*_ and *λ*_*i*_ using standard numerical methods. In the case where all boundary conditions are known, (1) is solved for all nodal pressures.

### Continuum blood flow model

Our hybrid model incorporates Poiseuille’s law to describe blood flow through arteriolar and venular branching structures, coupled to a volume-averaged description of blood velocity, **u** and pressure, *p*, within the capillary continuum domain ^47^ (see Figure 1C). The coupling occurs via points sources of flux located at the terminal branches of the discrete vasculature and feeding into the continuum capillary domain.

In the imaging of microvascular networks, limitations in field-of-view can lead to artificial discontinuities at the spatial boundary of the imaged network, which do not exist *in vivo*. In reality, blood flow continues from the imaged to the unimaged regions and vice versa at these artificial boundaries. To mitigate the impact of these imaging artefacts on our mathematical model, we introduce a *capillary network exchange term*, denoted as *β*(*p* − *p*_*c*_), where *p* represents the continuum pressure and *p*_*c*_ is the far-field capillary pressure. The rate of exchange, *β*, is not known *a priori*.

The exchange term is designed to facilitate capillary blood flow at the network boundary, thereby allowing for an interaction between the imaged capillary continuum and the adjacent, unimaged capillary network. It effectively relaxes the mass conservation constraint that would otherwise be imposed solely on the imaged discrete branching arteriolar and venular structures. In our model, the capillary network exchange term is applied throughout the domain and is essential for accurately representing the continuous flow of blood across the entire network, ensuring that the model accounts for interactions beyond the limited imaged area. Conservation of mass in the continuum domain is therefore defined as:

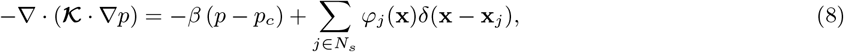

where ***𝒦*** is the hydraulic conductivity of the capillary network, *p*_*c*_ is a constant far-field pressure, *β* is the rate of capillary fluid drainage into neighbouring capillary network, *N*_*s*_ is the set of sources of flux into the continuum domain, and **x**_*j*_ and *φ*_*j*_ are the spatial coordinates and strength of source *j*, respectively. Source strengths are determined iteratively using (32) using a procedure outline in the proceeding sections.

Equation (8) is subject to the condition that

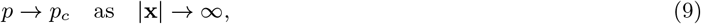

whereby it is assumed that the continuum pressure tends to a constant value, *p*_*c*_.

Hydraulic conductivity, ***𝒦***, is a 3×3 anisotropic tensor where we assume *𝒦*_*ij*_ *≪ 𝒦*_*ii*_ for *i, j* = 1, 2, 3 for *i≠j* ^23,28^. By setting *𝒦*_11_ = *κ, 𝒦*_22_ = *κa*^2^ and *𝒦*_33_ = *κb*^2^ we can introduce the linear transformation 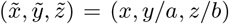 which reduces (8) to

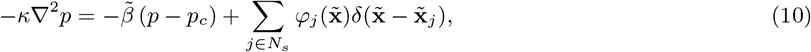

where 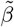 is the network boundary flow rate, *β*, in the scaled domain and (10) is subject to 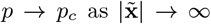 Whilst we observe later that the medulla/pons has a near isotropic hydraulic conductivity, which we provide this more generalisable linear scaling to enable others to apply our hybrid model to microvascular networks which exhibit a more anisotropic hydraulic conductivity tensor.

The solution to (10) is given by

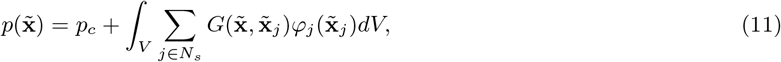

where *V* the volume of the tissue and 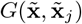 is a Green’s function which can be found by solving the adjoint problem:

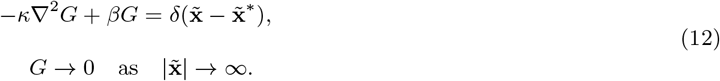

Here, 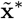 is a specific coordinate position.

Seeking a 3D, analytical representation of (12), we approximate 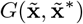 by distributing the delta function uniformly over a sphere of finite radius, *r*_0_. Considering a single source with local axisymmetric fluid flux, an axisymmetric spherical coordinate system is used whereby *r* = 0 corresponds to the source centre. Consequently, we seek the Green’s function solution, *G*(*r*), for the system

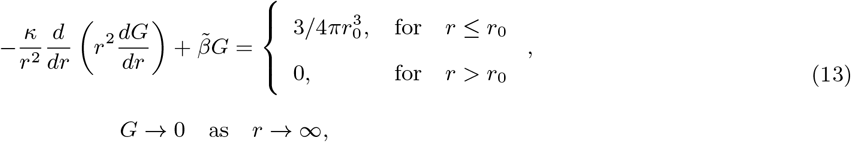

where 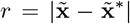 and the factor 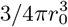 ensures unity when integrating 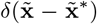 over the source volume. By ensuring continuity of *G*(*r*) and *G*^*′*^(*r*) at the interface *r* = *r*_0_, and that *G* is finite in the limit *r →* 0, we arrive at the solution

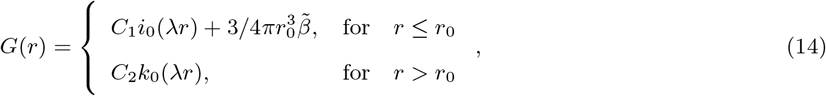

where *i*_0_ and *k*_0_ are the zeroth-order modified spherical Bessel functions of the first and second kinds, respectively,

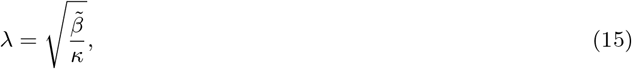

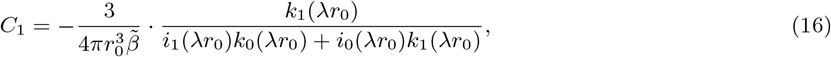

and

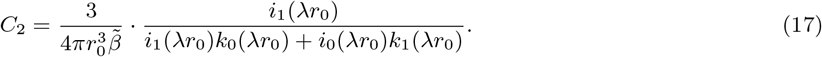

Figure 3 displays the behaviour of (14) for increasing values of the source radius, *r*_0_, and model parameter *λ* (see Appendix C for details on the limiting behaviour of *G* as *r*_0_ *→* 0). Increasing *r*_0_ descreases the amplitude of *G* yet broadens the function over *r*. In comparison, increasing *λ* both elevates the amplitude and value of *G* across *r*.

**Figure 3:**
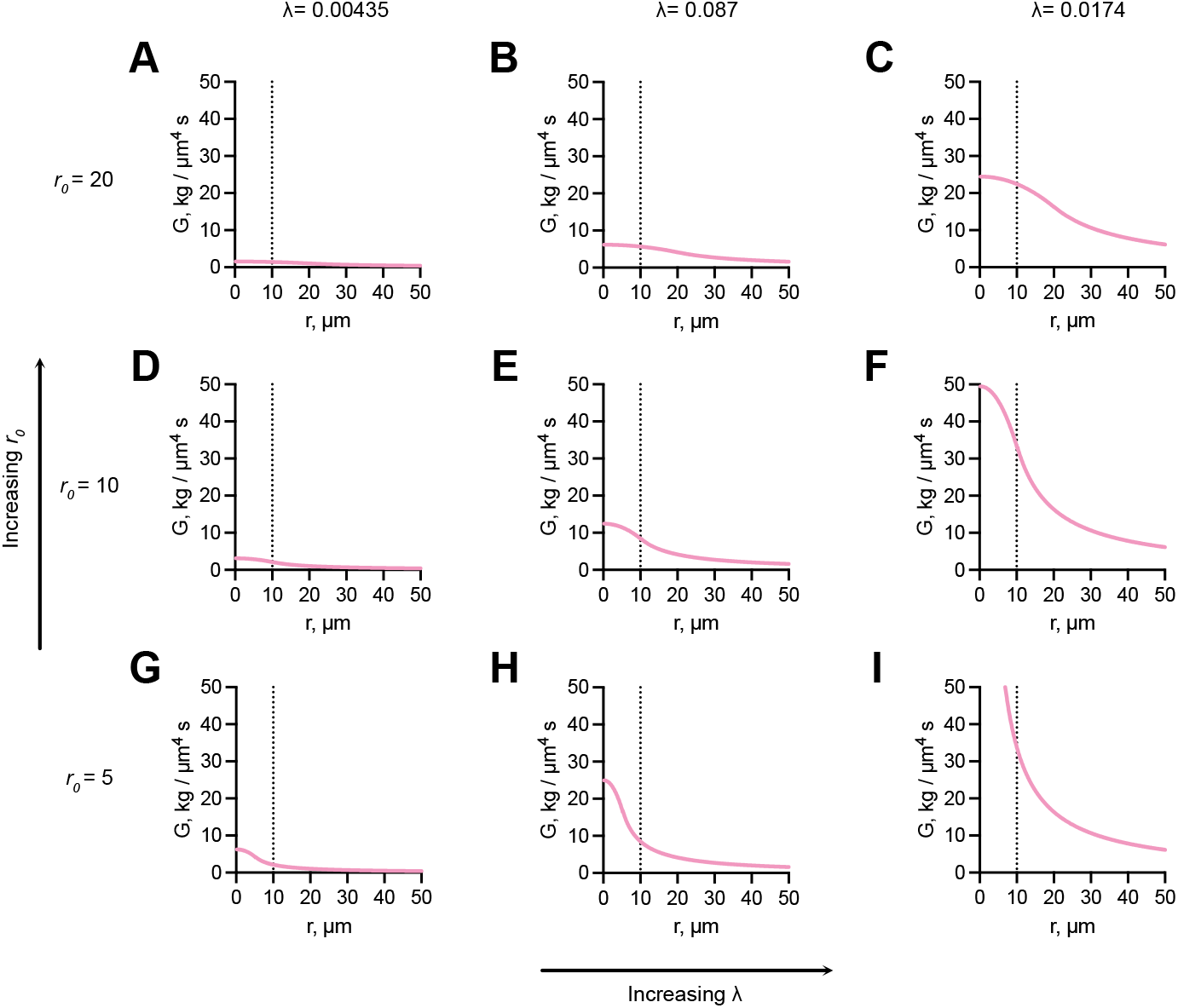
Behaviour of the Green’s function, *G*, with respect to source radius, *r*_0_, and parameter *λ*. The Green’s function, (14), is plotted for increasing values of *r*_0_ (bottom to top) and *λ* (left to right) where each increase corresponds to a doubling of the prior value. (B, E, H) Displays *λ* value used for medulla/pons baseline simulation (see Table 2).

We note that if blood pressure is modelled in an isolated tissue volume where flow is conserved between the branching structures, *β* = 0 and so (13) reduces to the model of Sweeney et al. ^17^ for vascular and interstitial fluid transport. See Appendix B for further details.

**Table 2:**
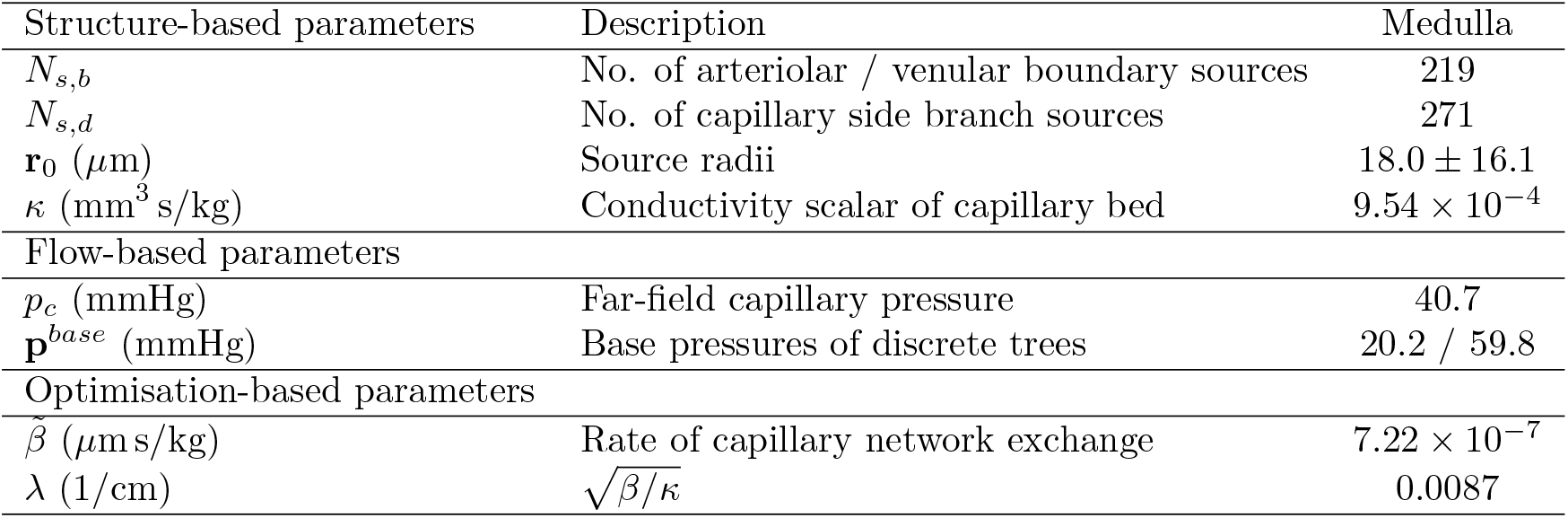
A summary of the parameter descriptions for the 3D discrete-continuum model and corresponding values for the medulla. Structural and flow-based parameters are assigned to calculate capillary continuum pressure. Optimisation-based parameters are fine-tuned to ensure flow conservation.

### Hydraulic conductivity of capillary tissue

Hydraulic conductivity is a tissue-specific parameter in our model which captures the resistance to flow in the capillary continuum domain. We calculate hydraulic conductivity by solving a micro-cell problem defined on 3D synthetic and periodic sub-units which are generated by sampling key features (such as diameters, lengths and connectivity) of the medulla capillary network. We average the micro-cell flow velocities, (18) to (28), on our synthetic, periodic architecture to calculate hydraulic conductivity tensors which describe how micro-scale flow properties are transmitted to the macro-scale of the medulla. First we provide a generalised description of solving the micro-cell problem in 3D, then our synthetic micro-cell generation method.

By assuming well-separated microscopic and macroscopic tissue length-scales, asymptotic homogenisation theory _*j*_ can be used to derive a system which averages microscopic capillary-scale flow to describe macroscopic tissue-scale fluid transport ^47^. An analytical solution to the multi-scale, micro-cell problem can be found by assuming non-leaky vessels, no-slip and no-flux at the capillary walls, which provides an expression for cell flux 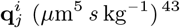:

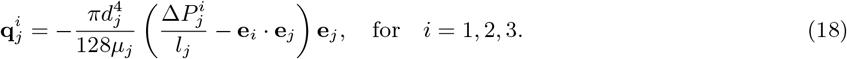

Here, *d*_*j*_, *l*_*j*_ and *µ*_*j*_ are the diameter (*µ*m), length (*µ*m) and blood viscosity (kg /*µ*m s, calculated empirically ^45^) for capillary *j*, respectively, 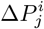 is micro-cell pressure gradient along the vessel length (*µ*m), **e**_*j*_ is the unit vector aligned with the capillary centreline and **e**_*i*_ is aligned with a tissue-scale principal direction. The principal axes can be defined based on tissue-scale features, for example, for the medulla/pons network the z-axis could be aligned with penetrating arteriolar / venular vessels. Micro-cell network architecture can then be generated with respect to these assigned axes.

Equation (18) is composed of a linear superposition of contributions proportional to the tissue-scale pressure gradient in each principal direction, **e**_*i*_. For vessel segments that are not aligned with the principal axes, we update **e**_*j*_ by decomposing the vessel segment’s orientation into its constituent vector components along the principal axes. The component weights of **e**_*j*_ are then adjusted so the forcing term **e**_*i*_ *·* **e**_*j*_ is correctly applied to the unaligned vessel.

The following details how a periodic boundary value problem is formed in terms of capillary-scale variations in flow, **q**^*i*^, and pressure, *P*^*i*^, over periodic micro-cells, enable the weightings of these contributions to be solved. Similar approaches have been used to calculate hydraulic conductivity for rat myocardium ^48^, human cerebral cortex ^23^ and rat mesentery ^35^.

Equation (18) can be rewritten in terms of micro-cell pressure at node *k*,

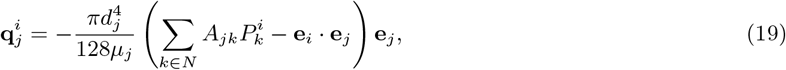

where 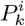 is the micro-cell pressure at node *k, N* is the set of all nodes and

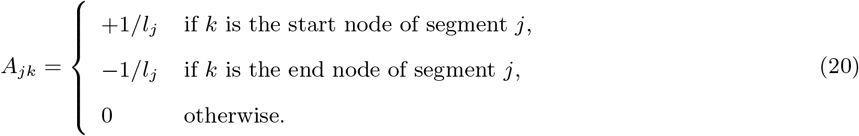

To solve for micro-cell flux and consequently micro-cell hydraulic conductivity, conservation of micro-cell flux, periodic Dirichlet boundary conditions for pressure and flux, and volume average of cell pressures equal to zero are enforced across the micro-cell.

Conservation of cell flux at nodes *m ∈ I*, where *I* is the set of interior nodes yields

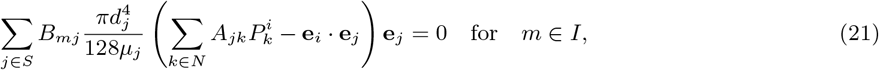

where *S* is the set of all segments and *B*_*mj*_ is defined as

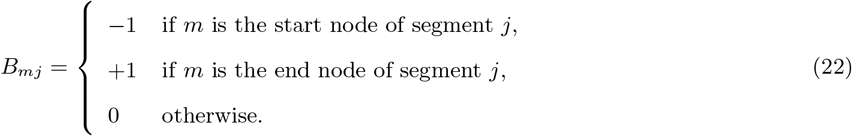

Separating boundary nodes into two sets, *B*_*in*_ and *B*_*out*_, where a node in *B*_*in*_ is paired with the node in *B*_*out*_ at the opposite side of the micro-cell, we impose periodic pressure boundary conditions via

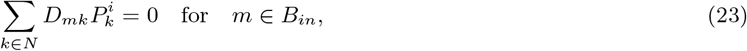

where

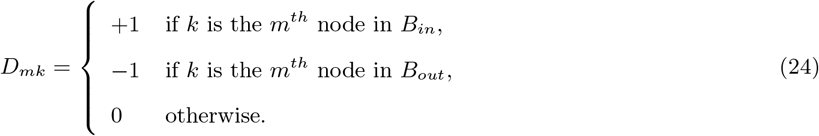

Similarly, for periodicity of flux, vessel flux leaving the *m*^*th*^ node of *B*_*out*_ is equal to the flux entering the *m*^*th*^ node of *B*_*in*_,

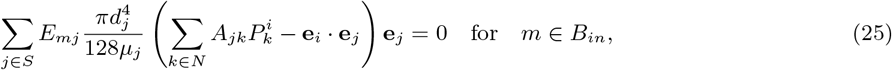

where

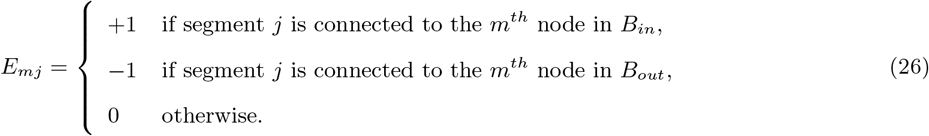

Finally, to ensure a unique solution, the volume average micro-cell pressure is defined to be zero:

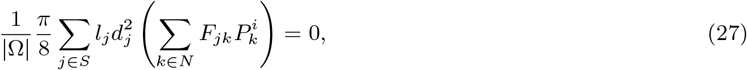

where

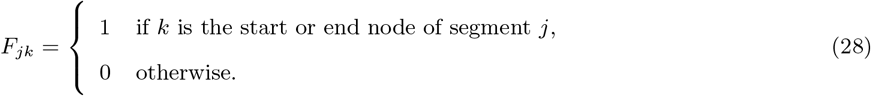

Equations (21), (23), (25) and (27) form three linear system of equations which can be solved for micro-cell nodal pressures *P*^*i*^ and consequently cell flux, (19). The hydraulic conductivity matrix ***𝒦*** = [***𝒦***^1^|***𝒦***^2^|***𝒦***^3^] (mm^3^s/kg) is then calculated via,

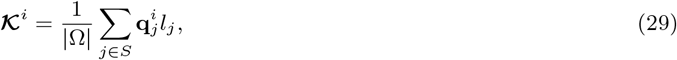

where Ω is the volume of the micro-cell and components 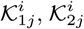 and 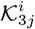 for *j* = 1, 2, 3 represent the volume-averaged flow in the **e**_1_, **e**_2_ and **e**_3_ directions, respectively, due to forcing in the **e**_*i*_-direction.

### Computational implementation of the discrete-continuum model

Here we outline the numerical implementation to predict the hydraulic conductivity of the capillary bed and capillary continuum pressure using our discrete-continuum model. A summary of the hybrid model, (6) to (17), is presented in Figure 4. In summary, boundary conditions are assigned to the bases and terminal nodes of the discrete branching structures. Each source to the continuum domain is iterated over by assigning a defined flow boundary condition and then pressures solved to enable the pressure-flow relationship across the discrete vasculature, via **M**^*net*^, to be characterised. In the capillary continuum component of our hybrid model, hydraulic conductivity of the domain is estimated via vascular homogenisation on a synthetically generated micro-cell, representative of the tissue-specific capillary architecture. Model parameters are assigned, ideally based on experimental or known literature values, and a Newton method is employed to solve for the capillary network exchange rate, *β*, which is minimised against a known net tissue flow. Once converged, capillary continuum pressures can be estimated using solved source fluxes.

**Figure 4:**
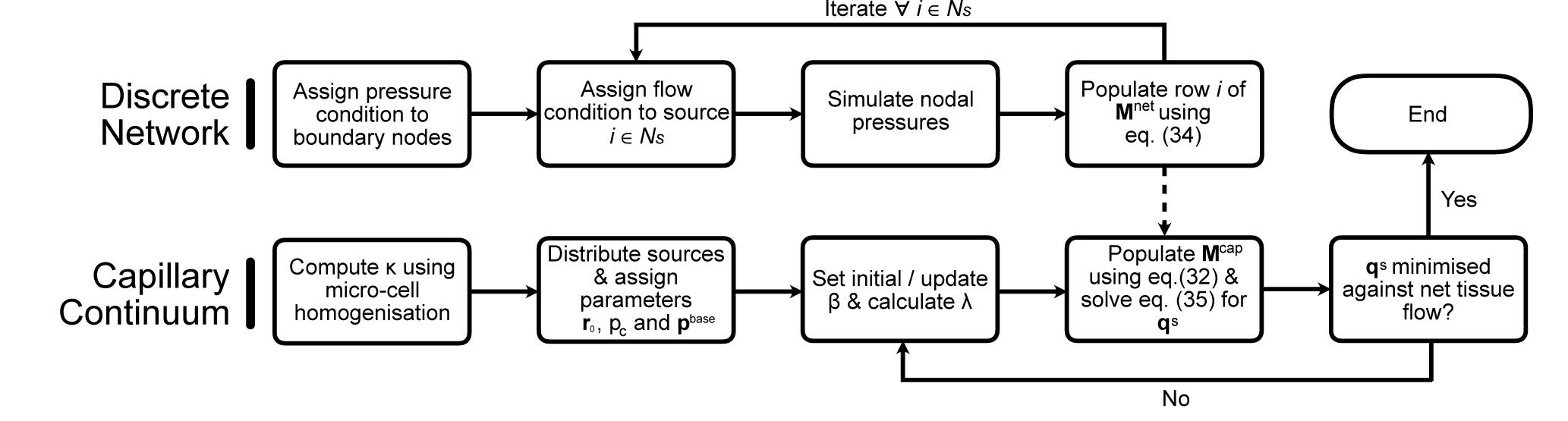
The discrete-continuum pipeline to solved for capillary bed pressure. (Top row) The discrete network matrix, **M**^*net*^, is populated by solving for discrete nodal pressures. (Bottom row) Capillary continuum source pressures found by solving the linear system, (34), (indicated by the dashed arrow). Conserving mass via optimisation of the capillary network exchange rate, *β*, ensures fluid flux by neighbouring capillary network is accurately calculated when validating against experimental, or in our case computational, methods.

### Generating 3D synthetic micro-cells

Key structural characteristics that influence perfusion in microvascular networks are vessel diameter and length, connectivity and vascular density (the network volume fraction with respect to tissue volume). Here, we define our methodology to generate 3D synthetic micro-cells which are representative of capillary vasculature by capturing these key attributes. For the medulla, we extract structural parameters of capillary networks directly from the fully-resolved network. In doing so, we approximate hydraulic conductivity for the medulla capillary bed which these sub-units represent. In the context of microvascular networks with largely unknown capillary networks, it is crucial to possess prior knowledge that allows users to create representative micro-cells. This prior knowledge can include known mean capillary diameters and lengths, along with experimental estimates of hydraulic conductivity. Having access to such information enables more informed predictions of hydraulic conductivity.

1. A 3D cuboidal grid lattice with dimensions 1×1×1 *µ*m^3^. We arbitrarily chose a lattice with 10×10 boundary nodes on each face of the lattice and 10*×*10*×*10 internal nodes, each of which interconnected by vessel segments (see Figure 1 and Appendix A).
2. To ensure physiological connectivity across the vessel lattice, and consequently physiological tissue perfusion, vessels were randomly removed until all nodes have a maximum of three vessel connections. Boundary vessels were exempt from removal to ensure boundary periodicity is maintained.
3. Vessels aligned with each axis were uniformly stretched in each dimension by factors randomly sampled from a log-normal distribution with mean and standard deviation for capillary lengths obtained from the medulla network. Similarly, diameters were individually sampled from known mean and standard deviation values.
4. Medulla vascular density was calculated to be 3.1%, and so if micro-cell vascular density is outside the interval 3.0 *±* 0.5%, step (3) is repeated.

Micro-cell hydraulic conductivity for a defined geometry is calculated using (29). Given micro-cell network generation is stochastic, steps (3) and (4) were repeated for a total of 10^3^ iterations, and average tensor components calculated.

We note that applying our micro-cell method to estimate hydraulic conductivity necessitates a prior understanding of the capillary structure. However, in practise, this information might not always be accessible. Under these circumstances, experimental approaches like dynamic contrast-enhanced MRI or photoacoustic imaging can be employed as effective alternatives. These methods enable the approximation of hydraulic conductivity through *in vivo* measurements of tissue perfusion and Darcy’s Law.

### Coupling discrete and continuum blood pressure

To calculate the continuum pressure field, we assume that the pressures at the inlet/outlet of the discrete arteriolar and venular trees, **p**^*base*^, and the far-field capillary pressure, *p*_*c*_ are known. The vector **p**^*base*^ is of length *N*_*s*_ where source *i* is assigned the base pressure of its respective vascular tree. From (11), the source pressure, 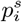, of source *i* at location 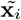 is given by

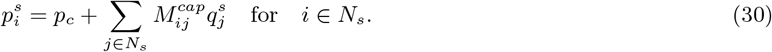

where

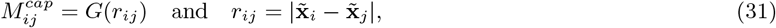

and 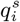 is the flow rate of source *i*.

The discrete blood flow model is used to calculate the pressures at nodes of the partial discrete networks as a function of the unknown flow conditions at the interface between the discrete and continuum domains:

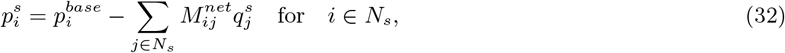

where the matrix **M**^*net*^ characterises the pressure-flow relationship across the discrete networks.

The set of source nodes includes capillary side branches of the vascular trees. Following Shipley et al. ^35^, in order for all source points to be terminal branches of their respective discrete network, dummy segments of 1 *µ*m length and diameter equal to the minimum diameter of its corresponding vascular tree were added. Then we perform a sequence of *i ∈ N*_*s*_ discrete flow calculations to populate 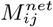 using (1)-(4). In each case, the following boundary conditions are applied: 1) if a node is a boundary node for the fully-discrete network (i.e., at the tissue boundary) a no flow boundary condition is applied; 2) the flow at source node *j* when *i* = *j* is set to *±*1 nl/min, where the positive / negative sign indicates whether the source belongs to an arteriole of venule, respectively, and zero for *i ≠ j*, so that (30) gives

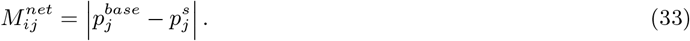

Combining the pressure solutions for (30) and (32) yields

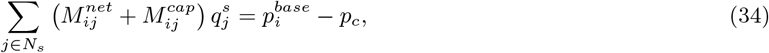

which can be solved for source strengths, **q**^*s*^.

### Boundary Conditions & Parameter Values

No flow or pressure information paired with the medulla dataset was obtained to parameterise either the fully-discrete or our hybrid model. Consequently, pressure boundary conditions at the bases of the arteriolar and venular trees were set using simulated cortex pressure-diameter curves from the literature ^9,10^. Vessel haematocrit was uniformly set to a value of 0.45. Based on previous studies, target blood pressure, **p**_0_, and vessel wall shear stress, *τ*_0_ were uniformly set to 41.3 mmHg and 5 dyn/cm^2^, respectively. Target blood pressure was calculate as the mean pressure between assigned arteriolar and venular boundary pressures, whereas target vessel was shear stress was assigned based on a prior study ^41^ as this data is not available from the medulla/pons. In comparison to boundary condition assignment, it has previously been shown that the model is not sensitive to assignment of these target values ^11^. Following the Fry et al. ^41^ flow optimisation scheme, *k*_*p*_ was set to a value of 0.1 and a final *k*_*τ*_ value of 7.21×10^12^ was found.

Parameter descriptions for the discrete-continuum model alongside values for the medulla network are provided in Table 2. In applying our hybrid model we found that to predict a physiological blood pressure distribution in the capillary continuum, the source radii need to be set relatively large compared to those in Shipley et al. ^35^ . As can be observed in the following analysis, the capillary pressure field tends to the far-field capillary pressure, *p*_*c*_, as the radii decrease. Consequently, we set source radii to the half-distance to the nearest neighbouring source so all sources act independently when solving for continuum pressure.

Base pressures, **p**^*base*^, were assigned using the simulated cortex pressure-diameter curves ^9,10^ and non-source, terminal boundaries were given no flow conditions. The far-field capillary pressure, *p*_*c*_, was assigned pragmatically using mean capillary blood pressure from the fully discrete network pressure solution and fixed at a value of 40.2 mmHg. However, in applications of the model, this can be assigned using an experimentally measure pressure value for the tissue of interest. Sensitivity analysis to test the influence of **r**_0_ and *p*_*c*_ on model predictions are also performed by increasing and decreasing thresholds.

With a predicted value for *κ*, a root-finding algorithm was used to compute *β*, and hence *λ*, by minimising the difference between the sum of source fluxes from the fully-discrete and hybrid pressure solutions (see Appendix D for further details). The fully-discrete pressure solution was used pragmatically as no experimental flow information was available for our medulla/pons network. Experimental methods which measure perfusion, such as arterial spin-labelling MRI, could be used calculate *β*.

Once source fluxes in the hybrid model are computed, pressure contour plots can be generated by computing (32) at grid points in the untransformed (*x, y, z*) domain. To enable comparison between the fully-discrete flow solution, we use the hybrid solution to predict nodal pressures at the spatial location of all nodes in the fully-discrete network.

All simulations were performed on a Macbook Pro with a 2.6 GHz 6-Core Intel Core i7 CPU and 16GB of RAM. Run-times for the our hybrid model and the flow estimation approach of ^41^ was < 1 min and < 10 min, respectively. Each model required less than 2 GB of RAM.

## Results

### Estimating hydraulic conductivity of the medulla capillary bed

A total of 10^3^ synthetic vascular micro-cells, characteristic of capillary networks in the medulla, were generated. For each periodic sub-unit, blood pressure was simulated (see Figure 5A-C) and hydraulic conductivity calculated using (29). Our computed averaged hydraulic conductivity tensor is given by

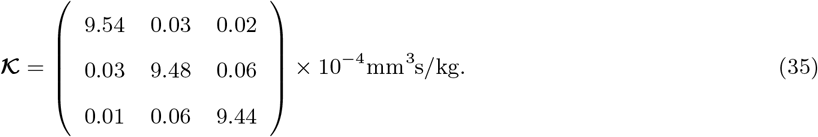

The diagonal components of the tensor are larger than the off-diagonal components thereby satisfying our assumption which enabled us to linearly transform the capillary continuum in (10). By introducing the linear scalings *a* = 0.997 and *b* = 0.995 where 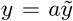 and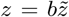, we reduce hydraulic conductivity to a scalar *κ* = 9.54×10^−4^ mm^3^s/kg in the transformed coordinate system. We note that the scalings *a ≈ b ≈* 1 is expected due to the stochastic method used to generate our synthetic micro-cells. These results is consistent with previous estimates of hydraulic conductivity for the myocardium ^48^ and cortex ^23^.

**Figure 5:**
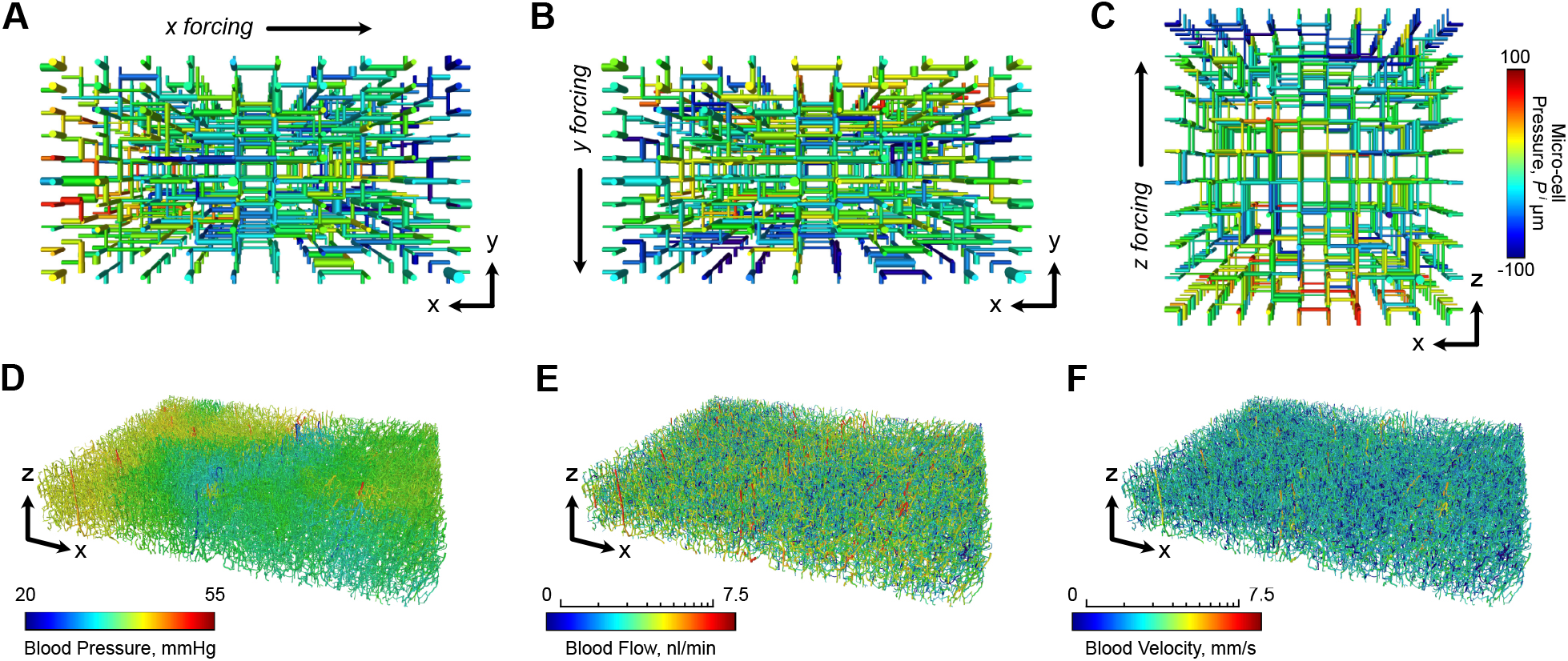
Micro-cell and discrete flow solutions. (A, B, C) Exemplar simulated micro-cell pressures for each principal direction *i* = 1, 2, 3 for *x, y* and *z*, respectively, used to estimated capillary hydraulic conductivity. Blood flow is simulated through the medulla using the fully-discrete model with predicted distributions of blood (D) pressure, (E) flow and (F) absolute velocity shown. Note, distributions in (E) and (F) are shown in a log scale.

### Discrete, complete network predictions

Arteriolar, venular and capillary pressures were predicted to be 44.63 *±* 5.04 mmHg, 31.37 *±* 5.34 mmHg and 40.70 *±* 3.03 mmHg, respectively (mean *±* standard deviation - see Figure 5D-F). To our knowledge, no experimental or computational predictions of microcirculatory blood pressure in the mouse medulla are available; however, our simulations are consistent with prior studies of the cortex ^9–12^. Additionally, minimum and maximum capillary pressures are within tolerances expected based on arteriolar and venular boundary condition assignment (see Table 3).

**Table 3:** Predicted pressures (mean *±* standard deviation) for the discrete and hybrid models applied to the medulla network.

| Simulated Pressures (mmHg) | Discrete | Hybrid |
| --- | --- | --- |
| Arteriolar Pressure | $44.63 \pm 5.04$ | - |
| Venular Pressure | $31.37 \pm 5.34$ | - |
| Capillary Pressure | $40.70 \pm 3.03$ | $40.09 \pm 2.035$ |
| Min. / Max. Capillary Pressure | 20.88 / 56.47 | 19.32 / 55.65 |
| Arteriolar Source Pressure | - | $44.40 \pm 3.41$ |
| Venular Source Pressure | - | $33.80 \pm 3.03$ |

### Hybrid modelling of blood pressure in the medulla

Continuum predictions of capillary blood pressure in the medulla were simulated using the parameters given by Table 2. With a hydraulic conductivity of *κ* = 9.54×10^−4^ mm^3^s/kg, the Green’s parameter *λ* was optimised to a value of 0.0087 cm ^−1^ with a rate of capillary network exchange of *β* = 7.22×10^−7^ *µ*m *s* kg^−1^, indicating very low levels of vessel flux across the tissue boundaries. The observed low levels of vessel flux across the capillary network boundaries in our simulations is physiologically reasonable on the basis that capillary blood flow and pressure is tightly controlled within the brain due to blood flow regulatory mechanisms and the spatially location of penetrating arterioles and venules, and blood flow regulatory mechanisms. These vessels play a significant role in directing blood supply to specific regions, ensuring efficient distribution throughout the brain which may minimise blood flow across the artificial capillary network boundaries.

The hybrid model calculated blood pressure at sources of 44.4 *±* 3.41 mmHg and 33.8 *±* 3.03 mmHg, for arterioles and venules, respectively, which are consistent with values calculated from the discrete model (see Table 2). Further, the hybrid and discrete model blood pressure predictions were strongly correlated for both arteriolar (Spearman correlation: *r* = 0.950 with *P* < 0.0001) and venular (Spearman correlation: *r* = 0.891 with *P* < 0.0001) sources, with pressures weighted towards the far-field capillary pressure, *p*_*c*_ = 40.7 mmHg. Compared to the fully-discrete predictions, the pressure difference at arteriolar sources increased particularly for those with pressure smaller than *p*_*c*_ (see Figure 6A). In comparison, venular source pressures exhibited increasingly larger errors for nodes with decreasingly lower pressures computed in the discrete model (see Figure 6B).

**Figure 6:**
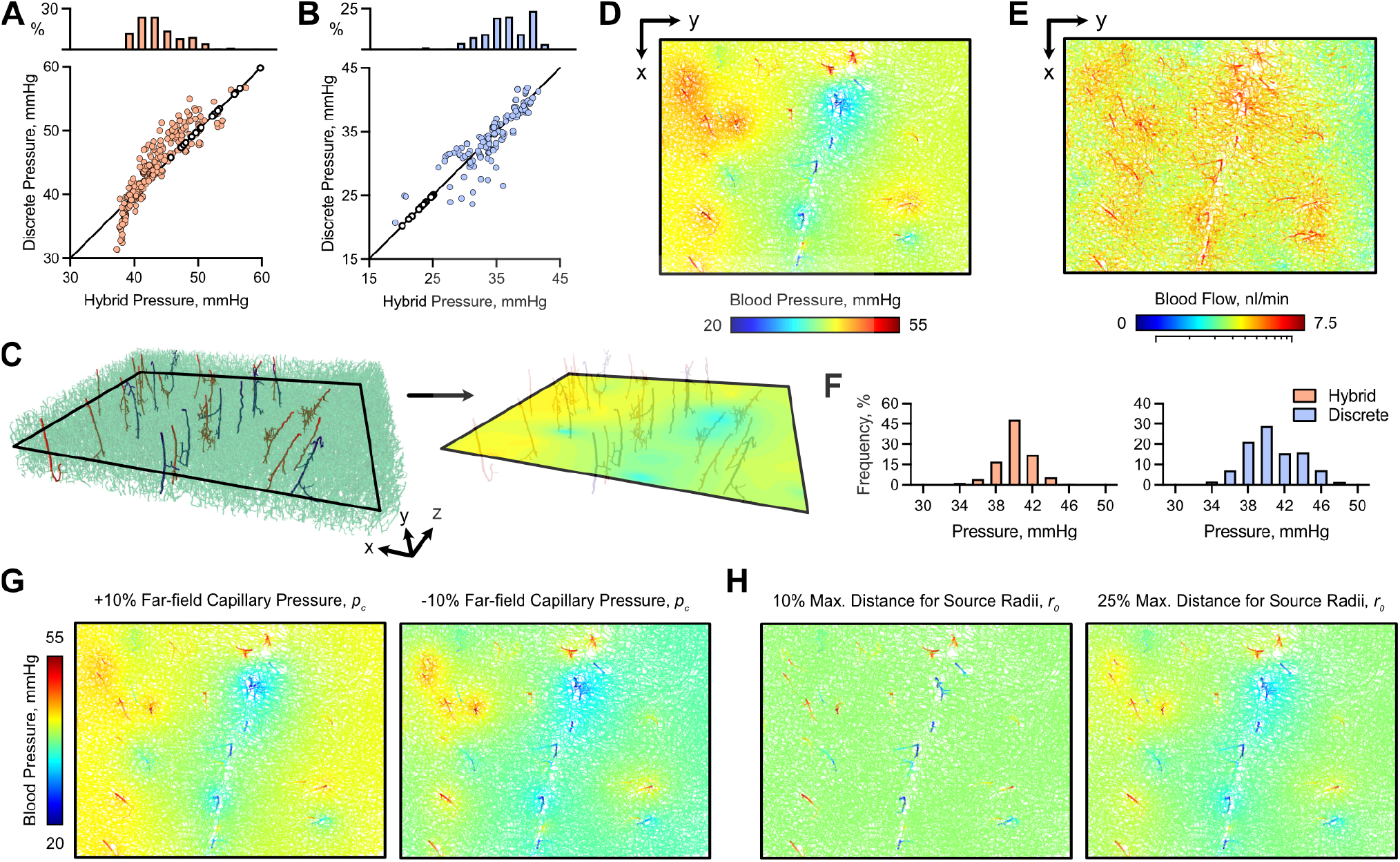
Hybrid blood pressure predictions in the mouse medulla. Discrete versus hybrid blood pressure predictions at (A) arteriolar and (B) venular point source locations with boundary conditions for each group and model shown in white. Distributions of hybrid source pressures for each vessel type are also illustrated. (C) Discrete-continuum model predictions of blood pressure in a 2D slice through the medulla. Here, arterioles, venules and capillaries are shown in red, blue and green, respectively. (D) A 2D intensity projection of the fully-resolved medulla network with continuum blood pressure predictions mapped onto the discrete vessels. (E) Associated absolute blood flow predictions. (F) Frequency distributions of blood pressures in the discrete and hybrid models at nodal locations in the fully-discrete medulla network. 2D intensity projections of the continuum blood pressure predictions mapped onto the fully-resolved medulla for (G) far-field capillary pressure, *p*_*c*_, and (H) source radii, *r*_0_, sensitivity analyses.

Once source pressures were computed, blood pressure was estimated across the entire capillary continua (see Figure 6C) which allowed our hybrid model estimates of blood pressure to be mapped onto the fully-discrete network (see Figure 6D,E). This enabled a comparison between blood pressures at all nodal locations of the fully-discrete network between the hybrid and discrete models. Capillary nodal pressures were found to be significantly different between the two models (Wilcoxon test: P< 0.0001) this could be due to the heavy weighting towards the far-field capillary pressure in the hybrid model (see Figure 6F). However, the relative error of segment pressure across the complete vascular network was 3.58 *±* 3.32% between the hybrid and discrete models. The relative flow segment error across the complete network was 177.3 *±* 408.2% which is likely due to the hybrid model providing an averaged, macro-scale prediction of capillary pressure and flow. This is compounded by the use of radially symmetric sources which do not capture the nuances of capillary tortuosity. Thus, errors at the micro-, vessel-scale are expected to be large.

### Blood pressure sensitivity to hybrid model parameters

Sensitivity analyses of hybrid model predictions were performed to assess the impact of key parameters. We focus on source pressures as an indicative metric. Unless stated otherwise, statistical significance (*P* < 0.05) compared to the baseline hybrid solution is calculated using a Kruskal-Wallis test.

Initially, we evaluated the sensitivity of the simulated source pressures to an arbitrary *±*10% fluctuation in the magnitude of the hydraulic conductivity, *κ*. This specific range was selected in alignment with the findings of El-Bouri and Payne ^23^, where hydraulic conductivity in the cortex was estimated using an alternative homogenization technique. In their study, a variability of *±*10% was observed across different micro-cell sizes. Considering the tightly regulated nature of blood flow in the brain, this variability range was considered suitable for application to the medulla / pons.

No significant difference was found for either arteriolar or venular sources when modifying *κ*, which indicates that the model is not susceptible to subtle changes in hydraulic conductivity (see Figure 7A). However, we noted that during testing, order of magnitude differences can significantly impact the simulated pressure distribution in the capillary bed which can result in a homogeneous or unphysiological pressure distributions.

**Figure 7:**
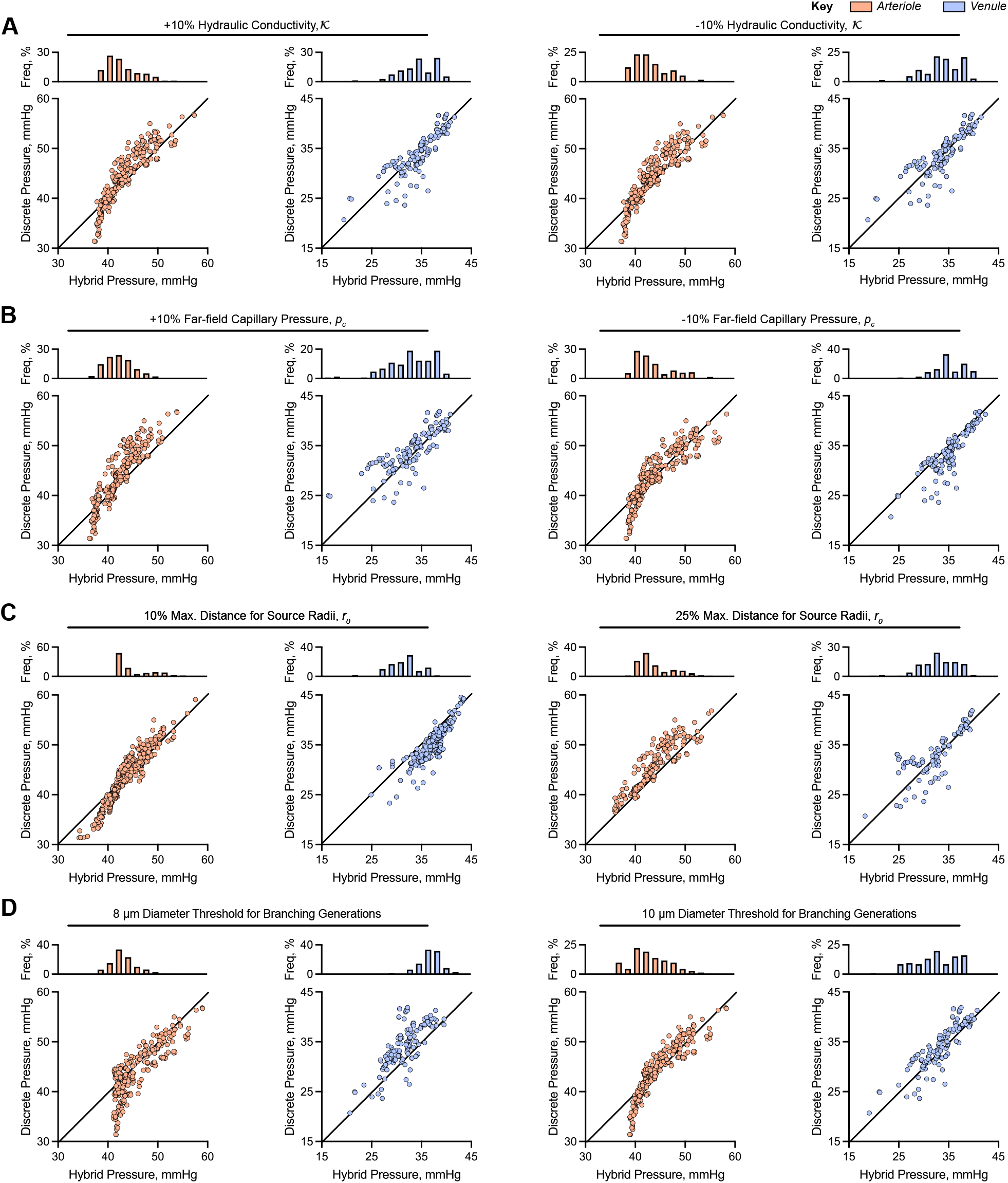
Sensitivity analysis to the hybrid model parameters. Source pressures solved using the hybrid model are plotted against nodal pressures at equivalent locations solved using the discrete model, with inset hybrid source pressure distributions. (A) Sensitivity to hydraulic conductivity scalar, *κ*, for a 10% (left) increase or (decrease) in magnitude. (B) Sensitivity to the far-field pressure, *p*_*c*_ for a 10% (left) increase or (decrease) in magnitude. (C) Sensitivity to the source radii **r**_0_ where each radius is set to (left) 10% or (right) 25% of the maximum distance to its nearest neighbour. (D) Sensitivity to the branching generation diameter threshold. Discrete branching vessels with a minimum diameter of (left) 8 *µ*m and 10 *µ*m are tested. Note, arterioles are shown in red and venules in blue.

Next, we perturb the far-field capillary pressure, *p*_*c*_, by *±*10% (see Figure 6G). Increasing *p*_*c*_ resulted in a significant decrease in venular pressures whereas no significant change was observed for arteriolar sources (see Figure 7B). In comparison, decreasing *p*_*c*_ widened the range of arteriolar source pressures, whereas venular sources exhibited a significant increase in simulated pressures. This is a consequence of optimising the capillary network exchange rate, *β*, to satisfy a known tissue flux, subject to the pressure range dictated by the pressures at the bases of the arteriolar and venular trees. In both cases, *β*-optimisation terminated outside of defined tolerances, with flow errors of 328.1 nl/min and −377.9 nl/min for +10% and −10% changes, respectively. This indicates that this parameter needs to be carefully parameterised in order to the reduce the relative conservation of mass error compared to the fully-discrete pressure solution.

For the baseline hybrid pressure (pre-sensitivity analysis) solution, each source radius was set equal to the half-distance to its closest neighbour. We found that decreasing these radii resulted in an unintuitive response to *β*. For example, distances of 25% and 10% relative to the closest neighbour, resulted in *β* values of 1.20×10^−7^ *µ*m *s* kg^−1^ and 0.98×10^3^ mm *s* kg^−1^. For the latter case arteriolar and venular source pressures exhibited a significant shift towards *p*_*c*_ = 40.7 mmHg (see Figure 6H). This may indicate that radii passed a critical threshold whereby the capillary capillary network exchange rate needed to compensate for the lack of flow transfer between arterioles and venules, due to large decreases in the pressure gradient between neighbouring sources (see Figure 7C). In comparison, for radii set to the quarter-distance, arteriolar source pressures shifted towards the far-field pressure. However, a response of similar magnitude was not observed for the venules, which may indicate that arterioles are be more susceptible to this parameter due to larger pressure gradients on the arteriolar side of the vascular hierarchy.

The final parameter we investigated was the minimum diameter threshold which was used to define the minimal cut-off of discrete vascular tree branches. Increasing the threshold from 9 *µ*m to 10 *µ*m resulted in a a non-significant change in arteriolar source pressures. However, venular source pressures exhibited a significant widening in distribution (see Figure 7D). By comparison setting the threshold to 8 *µ*m resulted in significant shifts in both the arteriolar (*P* < 0.05) and venular (*P* < 0.0001) pressures. In the case of arterioles, minimum source pressure was approximately equal to *p*_*c*_ which may be expected, as lowering the threshold increases correlation with the fully-discrete solution by allowing the hybrid model to incorporate more architectural heterogeneity. This would enable the discrete arteriolar networks to contain a greater portion of the pressure gradient. Comparatively, the change in venular distributions may be a product of a decrease in proximity to arteriolar sources as a result of a greater number of discrete branching orders. This may allow for lower venular source pressures at some locations.

## Discussion

The simultaneous measurement of blood pressure, flow and vascular structure *in vivo* is practically infeasible throughout whole tissues down to the microvascular resolution. This therefore motivates the development of mathematical models that can take vascular structures derived from biomedical images as inputs, predict fluid transport, and provide biological insights that are otherwise inaccessible. Several micron-scale imaging modalities exist which can extract complete vascular networks; however, the increasing size of these networks presents computational challenges for standard blood flow models. Moreover, some imaging modalities fail to retrieve the full detail of networks, particularly at the capillary scale, which limits applicability to standard discrete mathematical models.

Motivated by these limitations, we developed a hybrid discrete-continuum model to simulate blood pressure across incomplete 3D microvascular networks, extending upon prior 2D hybrid models ^29,35^. Our multiscale method models 1D Poiseuille flow through discrete branching arteriolar and venular structures, coupled to a single-phase Darcy model to estimate blood pressure through a continuum capillary bed. These discrete and continuum domains are connected via point sources of flux using an analytic Green’s function approach. Our model has two key tissue-specific parameters to effectively predict tissue blood pressure. Firstly, hydraulic conductivity in Darcy’s equation is approximated by local averaging of the capillary bed, by solving capillary flow through 3D, periodic and synthetically generated mico-cells ^21,29^. Secondly, conservation of mass between discrete branching structures is not imposed which allows for capillary flux at the tissue boundary. This rate of capillary flux is optimised against a known value of flow between the arteriolar and venular branches.

We apply our hybrid methodology to a complete vascular network from a mouse medulla, imaged using MF-HREM. Initially we simulated a baseline blood pressure solution with a standard 1D Poiseuille model across the fully-discrete medulla vasculature. Next, we retrospectively classified blood vessels in terms of arterioles, capillaries and venules which enabled us: 1) to quantify capillary structure in order to create medulla-specific micro-cells and predict the hydraulic conductivity of the medulla; 2) to provide discrete arteriolar and venular branching vessels for input into the hybrid model; and 3) an approximation of the capillary network exchange rate at the tissue boundaries. Using these data and parameter estimations, we simulated blood pressure across the capillary bed and mapped the continuum solution back onto the discrete capillary network to directly compare the hybrid model versus the fully-discrete blood pressure solution.

Our baseline hybrid capillary solution emulated pressure distributions observed in its fully-discrete counterpart. This demonstrates the effectiveness of the hybrid model to estimate physiological blood pressure distribution in the capillary bed in the absence of complete architectural data. Next we sought to understand how sensitive the model is to variance in a range of model parameters. In the first instance we targeted our estimation of medulla hydraulic conductivity. No experimental or computational value exists for the medulla in the literature, however, our initial approximation matched the order of magnitude for values found for other tissues ^23,48^. Artificially modifying hydraulic conductivity by *±*10% did not result in significant differences in capillary pressure. Nevertheless, tests of our model indicate order of magnitude differences can result in unphysiological blood pressure predictions.

Next, we altered the far-field capillary pressure and found that this is a key parameter when optimising for capillary network exchange, as incorrect assignment can significantly reduce the accuracy of flow conservation between arterioles and venules. In contrast, reducing the size of source radii still enabled capillary network exchange optimisation to converge; however, it caused local pressure gradients at each source to decrease. This results in a uniform pressure gradient across the capillary bed. Finally, we investigated the impact of retaining more structural arteriolar and venular architecture on the discrete-side of our model. By representing more vessels in a discrete fashion our hybrid model was better able to capture capillary pressure heterogeneity.

Here, we used the fully-discrete pressure solution as a gold standard to compare our hybrid model simulations against, despite it being an approximation. Nonetheless, our comparisons with the fully-discrete model provided insights into some limitations of the hybrid approach. The continuum representation of the capillary bed tended to smooth distributions towards the far-field capillary pressure. As a result, the hybrid model predicts lower levels of heterogeneity in pressure compared to the discrete model. This naturally presents several avenues to improve the modelling methodology. Firstly, the pressure field around each point source of flux into the continuum could be compartmentalised into several regions to better capture the pressure gradients in these regions ^28^. Secondly, constant values for the far-field capillary pressure and tissue hydraulic conductivity are an oversimplification, and so improved boundary conditions or spatially heterogeneous hydraulic conductivity could better approximate capillary pressure variability. Thirdly, capillary network exchange is parameterised by a flow balance calculated from the fully-discrete pressure solution. This is not practical in the experimental sense and so requires parameterisation via other means, such as perfusion MRI. Finally, red blood cell phase separation couple be incorporated into our hybrid model to provide a more realise prediction of haemodynamic features of the microcirculation. For the discrete component this is relatively straightforward; however, a continuum description would be more challenging due to the current mathematical formulation which prescribe periodic micro-cell boundary conditions.

## Conclusion

In summary, hybrid discrete-continuum models continue to present a promising approach to approximating the functionality of the microcirculation when limited structural information is available. Such models could prove complementary to *in vivo* experimentation which currently cannot observe tissue structure and transport across entire vascular networks, in particular for the smallest vessels.

## Acknowledgements

The authors also acknowledge support received from EPSRC (EP/L504889/1), Rosetrees Trust/Stoneygate Trust (M135-F1 and M601), the Wellcome Trust (WT100247MA) and Cancer Research UK (C44767/A29458).

## Author Contributions

Conceptualisation: PWS, RJS. Methodology: PWS, RJS. Software: PWS. Validation: PWS. Formal Analysis: PWS. Investigation: PWS. Resources: PWS, CW, SWS, RJS. Data Curation: PWS, CW. Writing - original draft: PWS, RJS. Writing - review & editing: PWS, CW, SWS, RJS. Visualisation: PWS. Supervision: RJS. Project Administration: PWS, RJS. Funding Acquisition: PWS, CW, SWS, RJS.

## Notes

### Competing Interest Statement

The authors have declared no competing interest.

### Summary of Updates

Edits to the content has been made throughout the manuscript.

## References

[1] Roger I Grant, David A Hartmann, Robert G Underly, Andrée Anne Berthiaume, Narayan R Bhat, and Andy Y Shih. Organizational hierarchy and structural diversity of microvascular pericytes in adult mouse cortex. Journal of Cerebral Blood Flow and Metabolism, pages 1–15, 2017. ISSN 15597016. doi: 10.1177/0271678X17732229.

[2] Timothy W Secomb. Hemodynamics. Comprehensive Physiology, 6(2):975–1003, 1 2016. ISSN 2040-4603. doi: 10.1002/cphy.c150038.

[3] T.W. Secomb, R. Hsu, M.W. Dewhirst, B. Klitzman, and J.F. Gross. Analysis of oxygen transport to tumor tissue by microvascular networks. International Journal of Radiation Oncology*Biology*Physics, 25(3):481–489, 2 1993. ISSN 03603016. doi: 10.1016/0360-3016(93)90070-C.

[4] Angela D’Esposito, Paul W. Sweeney, Morium Ali, Magdy Saleh, Rajiv Ramasawmy, Thomas A. Roberts, Giulia Agliardi, Adrien Desjardins, Mark F. Lythgoe, R. Barbara Pedley, Rebecca Shipley, and Simon Walker-Samuel. Computational fluid dynamics with imaging of cleared tissue and of in vivo perfusion predicts drug uptake and treatment responses in tumours. Nature Biomedical Engineering, 2(10):773–787, 10 2018. ISSN 2157-846X. doi: 10.1038/s41551-018-0306-y.

[5] Claire Walsh, Natalie A. Holroyd, Eoin Finnerty, Sean G. Ryan, Paul W. Sweeney, Rebecca J. Shipley, and Simon Walker-Samuel. Multifluorescence High-Resolution Episcopic Microscopy for 3D Imaging of Adult Murine Organs. Advanced Photonics Research, 2(10):2100110, 10 2021. ISSN 2699-9293. doi: 10.1002/ADPR.202100110.

[6] C. L. Walsh, P. Tafforeau, W. L. Wagner, D. J. Jafree, A. Bellier, C. Werlein, M. P. Kühnel, E. Boller, S. Walker-Samuel, J. L. Robertus, D. A. Long, J. Jacob, S. Marussi, E. Brown, N. Holroyd, D. D. Jonigk, M. Ackermann, and P. D. Lee. Imaging intact human organs with local resolution of cellular structures using hierarchical phase-contrast tomography. Nature Methods, 18(12):1532–1541, 11 2021. ISSN 15487105. doi: 10.1038/s41592-021-01317-x.

[7] Suhaas Anbazhakan, Pamela E. Rios Coronado, Ana Natalia L. Sy-Quia, Lek Wei Seow, Aubrey M. Hands, Mingming Zhao, Melody L. Dong, Martin R. Pfaller, Zhainib A. Amir, Brian C. Raftrey, Christopher K. Cook, Gaetano D’Amato, Xiaochen Fan, Ian M. Williams, Sawan K. Jha, Daniel Bernstein, Koen Nieman, Anca M. Pas ca, Alison L. Marsden, and Kristy Red Horse. Blood flow modeling reveals improved collateral artery performance during the regenerative period in mammalian hearts. Nature Cardiovascular Research 2022 1:8, 1 (8):775–790, 8 2022. ISSN 2731-0590. doi: 10.1038/s44161-022-00114-9.

[8] Maxime Berg, Natalie Holroyd, Claire Walsh, Hannah West, Simon Walker-Samuel, and Rebecca Shipley. Challenges and opportunities of integrating imaging and mathematical modelling to interrogate biological processes. International Journal of Biochemistry and Cell Biology, 146:106195, 5 2022. ISSN 18785875. doi: 10.1016/j.biocel.2022.106195.

[9] S. Lorthois, F. Cassot, and F. Lauwers. Simulation study of brain blood flow regulation by intra-cortical arterioles in an anatomically accurate large human vascular network. Part II: Flow variations induced by global or localized modifications of arteriolar diameters. NeuroImage, 54(4):2840–2853, 2011. ISSN 10538119. doi: 10.1016/j.neuroimage.2010.10.040.

[10] S. Lorthois, F. Cassot, and F. Lauwers. Simulation study of brain blood flow regulation by intra-cortical arterioles in an anatomically accurate large human vascular network: Part I: Methodology and baseline flow. NeuroImage, 54(2):1031–1042, 2011. ISSN 10538119. doi: 10.1016/j.neuroimage.2010.09.032.

[11] Paul W. Sweeney, Simon Walker-Samuel, and Rebecca J. Shipley. Insights into cerebral haemodynamics and oxygenation utilising in vivo mural cell imaging and mathematical modelling. Scientific Reports, 8(1):1373, 12 2018. ISSN 20452322. doi: 10.1038/s41598-017-19086-z.

[12] Jose T Celaya-Alcala, Grace V Lee, Amy F Smith, Bohan Li, Sava Sakadžić, David A Boas, and Timothy W Secomb. Simulation of oxygen transport and estimation of tissue perfusion in extensive microvascular networks: Application to cerebral cortex. Journal of Cerebral Blood Flow and Metabolism, 41(3):656–669, 6 2021. ISSN 15597016. doi: 10.1177/0271678X20927100.

[13] Timothy W. Secomb, Jonathan P. Alberding, Richard Hsu, Mark W. Dewhirst, and Axel R. Pries. Angiogenesis: An Adaptive Dynamic Biological Patterning Problem. PLoS Computational Biology, 9(3), 2013. ISSN 1553734X. doi: 10.1371/journal.pcbi.1002983.

[14] Jonathan P. Alberding and Timothy W. Secomb. Simulation of angiogenesis in three dimensions: Application to cerebral cortex. PLOS Computational Biology, 17(6):e1009164, 6 2021. ISSN 1553-7358. doi: 10.1371/JOURNAL.PCBI.1009164.

[15] Vasileios Vavourakis, Peter A. Wijeratne, Rebecca Shipley, Marilena Loizidou, Triantafyllos Stylianopoulos, and David J. Hawkes. A Validated Multiscale In-Silico Model for Mechano-sensitive Tumour Angiogenesis and Growth. PLoS Computational Biology, 13(1):e1005259, 1 2017. ISSN 15537358. doi: 10.1371/journal.pcbi.1005259.

[16] Vasileios Vavourakis, Triantafyllos Stylianopoulos, and Peter A. Wijeratne. In-silico dynamic analysis of cytotoxic drug administration to solid tumours: Effect of binding affinity and vessel permeability. PLoS Computational Biology, 14(10):e1006460, 10 2018. ISSN 1553-7358. doi: 10.1371/journal.pcbi.1006460.

[17] Paul W. Sweeney, Angela D’Esposito, Simon Walker-Samuel, and Rebecca J. Shipley. Modelling the transport of fluid through heterogeneous, whole tumours in silico. PLOS Computational Biology, 15(6):e1006751, 6 2019. ISSN 1553-7358. doi: 10.1371/journal.pcbi.1006751.

[18] Emma Brown, Joanna Brunker, and Sarah E Bohndiek. Photoacoustic imaging as a tool to probe the tumour microenvironment. Disease Models and Mechanisms, 12(7):dmm039636, 2019. ISSN 17548411. doi: 10.1242/dmm.039636.

[19] Emma L. Brown, Thierry L. Lefebvre, Paul W. Sweeney, Bernadette J. Stolz, Janek Gröhl, Lina Hacker, Ziqiang Huang, Dominique Laurent Couturier, Heather A. Harrington, Helen M. Byrne, and Sarah E. Bohndiek. Quantification of vascular networks in photoacoustic mesoscopy. Photoacoustics, 26:100357, 6 2022. ISSN 22135979. doi: 10.1016/j.pacs.2022.100357.

[20] S Jonathan Chapman, Rebecca J Shipley, and Rossa Jawad. Multiscale modeling of fluid transport in tumors. Bulletin of mathematical biology, 70(8):2334–57, 11 2008. ISSN 1522-9602. doi: 10.1007/s11538-008-9349-7.

[21] Rebecca J Shipley and S Jonathan Chapman. Multiscale modelling of fluid and drug transport in vascular tumours. Bulletin of mathematical biology, 72(6):1464–91, 8 2010. ISSN 1522-9602. doi: 10.1007/s11538-010-9504-9.

[22] Tiina Roose and Melody A. Swartz. Multiscale modeling of lymphatic drainage from tissues using homogenization theory. Journal of Biomechanics, 45(1):107–115, 1 2012. ISSN 0021-9290. doi: 10.1016/J.JBIOMECH.2011.09.015.

[23] Wahbi K El-Bouri and Stephen J Payne. Multi-scale homogenization of blood flow in 3-dimensional human cerebral microvascular networks. Journal of Theoretical Biology, 380:40–47, 9 2015. ISSN 10958541. doi: 10.1016/j.jtbi.2015.05.011.

[24] R. Penta and D. Ambrosi. The role of the microvascular tortuosity in tumor transport phenomena. Journal of Theoretical Biology, 364:80–97, 1 2015. ISSN 10958541. doi: 10.1016/j.jtbi.2014.08.007.

[25] R. Penta, D. Ambrosi, and A. Quarteroni. Multiscale homogenization for fluid and drug transport in vascularized malignant tissues. Mathematical Models and Methods in Applied Sciences, 25(1):79–108, 11 2015. ISSN 02182025. doi: 10.1142/S0218202515500037.

[26] Pietro Mascheroni and Raimondo Penta. The role of the microvascular network structure on diffusion and consumption of anticancer drugs. International Journal for Numerical Methods in Biomedical Engineering, 33 (10):e2857, 10 2017. ISSN 20407947. doi: 10.1002/cnm.2857.

[27] Rebecca J. Shipley, Paul W. Sweeney, Stephen J. Chapman, and Tiina Roose. A four-compartment multiscale model of fluid and drug distribution in vascular tumours. International Journal for Numerical Methods in Biomedical Engineering, 36(3):e3315, 3 2020. ISSN 2040-7947. doi: 10.1002/CNM.3315.

[28] Myriam Peyrounette, Yohan Davit, Michel Quintard, and Sylvie Lorthois. Multiscale modelling of blood flow in cerebral microcirculation: Details at capillary scale control accuracy at the level of the cortex. PLoS ONE, 13 (1):e0189474, 2018. doi: 10.1371/journal.pone.0189474.

[29] Amy F. Smith, Bianca Nitzsche, Martin Maibier, Axel R. Pries, and Timothy W. Secomb. Microvascular hemodynamics in the chick chorioallantoic membrane. Microcirculation, 23(7):512–522, 8 2016. ISSN 15498719. doi: 10.1111/micc.12301.

[30] M. Kojic, M. Milosevic, V. Simic, E. J. Koay, J. B. Fleming, S. Nizzero, N. Kojic, A. Ziemys, and M. Ferrari. A composite smeared finite element for mass transport in capillary systems and biological tissue. Computer methods in applied mechanics and engineering, 324:413–437, 9 2017. ISSN 0045-7825. doi: 10.1016/J.CMA.2017.06.019.

[31] S. J. Payne and W. K. El-Bouri. Modelling dynamic changes in blood flow and volume in the cerebral vasculature. NeuroImage, 176:124–137, 8 2018. ISSN 10959572. doi: 10.1016/j.neuroimage.2018.04.037.

[32] Wahbi K. El-Bouri and Stephen J. Payne. Investigating the effects of a penetrating vessel occlusion with a multi-scale microvasculature model of the human cerebral cortex. NeuroImage, 172:94–106, 5 2018. ISSN 10959572. doi: 10.1016/j.neuroimage.2018.01.049.

[33] Johannes Kremheller, Anh Tu Vuong, Bernhard A. Schrefler, and Wolfgang A. Wall. An approach for vascular tumor growth based on a hybrid embedded/homogenized treatment of the vasculature within a multiphase porous medium model. International Journal for Numerical Methods in Biomedical Engineering, 35(11):e3253, 11 2019. ISSN 2040-7947. doi: 10.1002/CNM.3253.

[34] Ettore Vidotto, Timo Koch, Tobias Koppl, Rainer Helmig, and Barbara Wohlmuth. Hybrid Models for Simulating Blood Flow in Microvascular Networks. Multiscale modeling & simulation : a SIAM interdisciplinary journal, 17(3):1076–1102, 9 2019. ISSN 15403467. doi: 10.1137/18M1228712.

[35] Rebecca J Shipley, Amy F Smith, Paul W Sweeney, Axel R Pries, and Timothy W Secomb. A hybrid discrete–continuum approach for modelling microcirculatory blood flow. Mathematical Medicine and Biology: A Journal of the IMA, 37(1):40–57, 3 2020. ISSN 1477-8599. doi: 10.1093/imammb/dqz006.

[36] Timo Koch, Martin Schneider, Rainer Helmig, and Patrick Jenny. Modeling tissue perfusion in terms of 1d-3d embedded mixed-dimension coupled problems with distributed sources. Journal of Computational Physics, 410: 109370, 6 2020. ISSN 0021-9991. doi: 10.1016/J.JCP.2020.109370.

[37] Johannes Kremheller, Sebastian Brandstaeter, Bernhard A. Schrefler, and Wolfgang A. Wall. Validation and parameter optimization of a hybrid embedded/homogenized solid tumor perfusion model. International Journal for Numerical Methods in Biomedical Engineering, 37(8):e3508, 8 2021. ISSN 20407947. doi: 10.1002/cnm.3508.

[38] L. J. Cooper, K. R. Daly, P. D. Hallett, M. Naveed, N. Koebernick, A. G. Bengough, T. S. George, and T. Roose. Fluid flow in porous media using image-based modelling to parametrize Richards’ equation. In Proceedings of the Royal Society A: Mathematical, Physical and Engineering Sciences, volume 473. The Royal Society Publishing, 11 2017. doi: 10.1098/rspa.2017.0178.

[39] K. R. Daly and T. Roose. Determination of macro-scale soil properties from pore-scale structures: Model derivation. Proceedings of the Royal Society A: Mathematical, Physical and Engineering Sciences, 474(2209), 2018. ISSN 14712946. doi: 10.1098/rspa.2017.0141.

[40] Benyi Xiong, Anan Li, Yang Lou, Shangbin Chen, Ben Long, Jie Peng, Zhongqin Yang, Tonghui Xu, Xiao-quan Yang, Xiangning Li, Tao Jiang, Qingming Luo, and Hui Gong. Precise cerebral vascular atlas in stereotaxic coordinates of whole mouse brain. Frontiers in Neuroanatomy, 11:128, 12 2017. ISSN 16625129. doi: 10.3389/fnana.2017.00128.

[41] Brendan C. Fry, Jack Lee, Nicolas P. Smith, and Timothy W. Secomb. Estimation of Blood Flow Rates in Large Microvascular Networks. Microcirculation, 19:530–538, 2012. ISSN 10739688. doi: 10.1111/j.1549-8719.2012.00184.x.

[42] Spyros k. Stamatelos, Eugene Kim, Arvind P. Pathak Popel, and Aleksander S. A bioimage informatics based reconstruction of breast tumor microvasculature with computational blood flow predictions. Microvascular Research, 91:8–21, 2014. doi: 10.1016/j.mvr.2013.12.003.

[43] Amy Smith. Multi-scale modelling of blood flow in the coronary microcirculation. PhD thesis, University of Oxford, 2013.

[44] Paul W. Sweeney. Realistic numerical image-based modelling of biological tissue substrates. PhD thesis, University College London, 2018. URL http://discovery.ucl.ac.uk/id/eprint/10049410.

[45] A R Pries and T W Secomb. Microvascular blood viscosity in vivo and the endothelial surface layer. American journal of physiology. Heart and circulatory physiology, 289(6):H2657–H2664, 2005. ISSN 0363-6135. doi: 10.1152/ajpheart.00297.2005.

[46] A R Pries, K Ley, M Claassen, and P Gaehtgens. Red cell distribution at microvascular bifurcations. Microvascular research, 38(1):81–101, 7 1989. ISSN 0026-2862.

[47] Rebecca J Shipley. Multiscale modelling of fluid and drug transport in vascular tumours. PhD thesis, University of Oxford, 2008.

[48] Amy F Smith, Rebecca J Shipley, Jack Lee, Gregory B Sands, Ian J LeGrice, and Nicolas P Smith. Transmural variation and anisotropy of microvascular flow conductivity in the rat myocardium. Annals of biomedical engineering, 42(9):1966–77, 9 2014. ISSN 1573-9686. doi: 10.1007/s10439-014-1028-2.

