## Supplementary Material for "A three-dimensional, discrete-continuum model of blood pressure in microvascular networks"

[*]Correspondance

#### Appendix A: Supplementary Figures

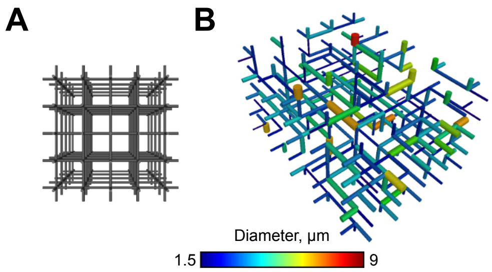


Figure S1: Generating 3D synthetic, periodic micro-cell. (A) Step 1: A 3D cuboidal grid lattice with dimensions $1 \times 1 \times 1$ $\mu$m$^3$ with, for example, $5 \times 5$ boundary nodes on each face of the cuboid. Each node is connected by a vessel segment (shown in grey). (B) Step 2 to 3: vessels are pruned to ensure physiological connectivity and the micro-cell is stretched in each axial direction based on known mean capillary lengths.

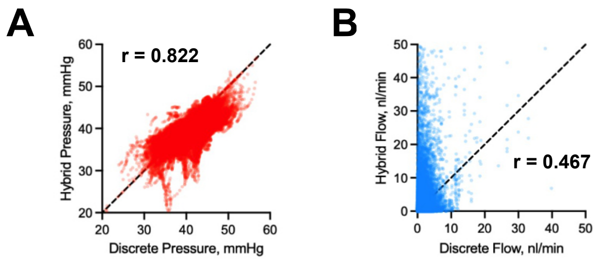


Figure S2: Plotting vessel segment (A) pressures (mmHg) and (B) absolute flow rates (nl/min) between predictions from the hybrid and fully discrete model. R-values were computed using Spearman's Correlation. Relative errors are $3.58 \pm 3.32\%$ and $177.3 \pm 408.2\%$ for segment pressure and flow, respectively.

### Appendix B: Flow conservation between branching vasculatures

Flow conservation between branching blood vessels results in $\beta = 0$ reducing (13) to

$$-\frac{\kappa}{r^2}\frac{d}{dr}\left(r^2\frac{dG}{dr}\right) = \begin{cases} 3/4\pi r_0^3, & \text{for} \quad r \le r_0 \\ 0, & \text{for} \quad r > r_0 \end{cases}, \qquad G \to 0 \quad \text{as} \quad r \to \infty. \tag{1}$$

This is equivalent to the model of Sweeney et al. [1] for interstitial fluid transport which has the solution

$$G(r) = \begin{cases} \dfrac{1}{8\pi\kappa r_0}\left[3 - \left(\dfrac{r}{r_0}\right)^2\right], & \text{for} \quad r \le r_0, \\ \dfrac{1}{4\pi\kappa r}, & \text{for} \quad r > r_0. \end{cases} \tag{2}$$

This solution can simply replace (14) in the hybrid model and since $\beta = 0$, removes the requirement to run the $\beta$-optimisation algorithm.

### Appendix C: The limit of the Green's function, $G$

Here, we analyse the limit of our Green's function, $G$, given by eq. (14) in the main manuscript as $r_0 \to 0$. We evaluate its limits using series expansions of the Bessel functions near $r_0 = 0$ as using L'Hôpital's rule leads to an indeterminate form.

The series expansions of the modified Spherical Bessel functions of the first kind, $i_0(r_0)$ and $i_1(r_0)$, at

$r_0 = 0$ for $r_0 > 0$ are:

$$i_0(r_0) = 1 + \frac{r_0^2}{6} + \frac{r_0^4}{120} + \dots \tag{3}$$

and

$$i_1(r_0) = \frac{r_0}{3} + \frac{r_0^3}{30} + \frac{r_0^5}{840} + \dots. \tag{4}$$

Similarly, the series expansion of the modified Bessel functions of the second kind, $k_0(r_0)$ and $k_1(r_0)$, at $r_0 = 0$ for $r_0 > 0$ are:

$$k_0(r_0) = \frac{1}{r_0} - 1 + \frac{r_0}{2} + \dots \tag{5}$$

and

$$k_1(r_0) = \frac{1}{r_0^2} - \frac{1}{2} + \frac{r_0}{3} + \dots. \tag{6}$$

Setting $r$ and $\lambda$ as constant values, we can evaluate the limit of $G$ at $r_0 = 0$:

$$\begin{aligned}
G(r) &= \frac{3}{4\pi\tilde{\beta}} \cdot \lim_{r_0 \to 0} \frac{1}{r_0^3} \left[ \frac{i_0(\lambda r) k_1(\lambda r_0)}{i_1(\lambda r_0) k_0(\lambda r_0) + i_0(\lambda r_0) k_1(\lambda r_0)} + 1 \right] \\
&= \frac{3}{4\pi\tilde{\beta}} \cdot \lim_{r_0 \to 0} \frac{1}{r_0^3} \\
&\cdot \left[ \frac{i_0(\lambda r)(\frac{1}{(\lambda r_0)^2} - \frac{1}{2} + \frac{\lambda r_0}{3} + \dots)}{(\frac{\lambda r_0}{3} + \frac{(\lambda r_0)^3}{30} + \frac{(\lambda r_0)^5}{840} + \dots)(\frac{1}{\lambda r_0} - 1 + \frac{\lambda r_0}{2} + \dots) + (1 + \frac{(\lambda r_0)^2}{6} + \frac{(\lambda r_0)^4}{120} + \dots)(\frac{1}{(\lambda r_0)^2} - \frac{1}{2} + \frac{\lambda r_0}{3} + \dots)} + 1 \right] \\
&= \frac{3}{4\pi\tilde{\beta}} \cdot \lim_{r_0 \to 0} \frac{1}{r_0^3} \left[ i_0(\lambda r) \cdot \frac{(\frac{1}{(\lambda r_0)^2} - \frac{1}{2} + \frac{\lambda r_0}{3} + \dots)}{(\frac{1}{(\lambda r_0)^2} - \frac{\lambda r_0}{3} - \frac{(\lambda r_0)^2}{20} - \frac{(\lambda r_0)^3}{30} + \dots)} + 1 \right] \\
&\to \infty \quad \text{as} \quad r_0 \to 0.
\end{aligned} \tag{7}$$

### Appendix D: An Analytical Solution to the Micro-Cell Problem

The following derivation was performed by Smith [2] and reported in Sweeney [3].

Shipley and Chapman [4] developed a cell problem which computes the macro-scale resistance to fluid transport by solving for flow through periodic micro-cells (given by (64) - (67) in the study). The cell problem is subject to the conditions that micro-cell flux is conserved, boundary pressures and flux are periodic, and the volume average of the cells pressures is zero.

Derivation of the Shipley and Chapman [4] model assumed that blood viscosity is constant and so the non-Newtonian behaviour of blood in the microcirculation can be described by the empirical viscosity laws of Pries and Secomb[5]. As such, in the case of impermeable capillaries, the cell problem reduces to

$$\nabla_X \cdot \mathbf{w}_j^i = 0 \quad \text{in} \quad \Omega_c \tag{8a}$$

$$\mathbf{e}_i = \nabla_X P_j^i - \mu_{abs} \nabla_X^2 \mathbf{w}_j^i \quad \text{in} \quad \Omega_c, \tag{8b}$$

$$\mathbf{w}_j^i = \mathbf{0} \quad \text{on} \quad \Gamma, \tag{8c}$$

where $\mathbf{w}_j^i$ and $P_j^i$ are the micro-cell velocity and blood pressure for capillary segment $j$ in each principal axis $i$. Equation (8) is also subject to the volume averaged periodic condition over all micro-cell capillaries,

$$\left\langle P^i \right\rangle = 0. \tag{9}$$

Recalling that the variable $\mathbf{X}$ represents the micro-cell length scale.

Using the scheme outlined by Shipley[6], Smith[2] sought an analytical approximation for $\mathbf{w}_j^i$ and $P_j^i$ to the system (8) which is analogous to Poiseuille flow but with periodic tissue-scale forcing. Capillaries were assumed to be thin, straight cylinders of constant circular cross-section. The following derivation considers a single capillary, and so the subscript $j$ is dropped for convenience. A local non-dimensional cylindrical coordinate system $(R, \theta, s)$ was used where $s$ is the coordinate aligned with the vessel centreline.

The variable $\delta$ was defined as the non-dimensional capillary radius which allowed the introduction of the stretched variable $r = R/\delta$. Here, $R$ is the dimensional radius of the vessel, along with the dimensionless re-scalings, $U$, $V$ and $W$ for the micro-cell velocity components, and so

$$\mathbf{w}^i = U w_r^i \mathbf{e}_r + V w_\theta^i \mathbf{e}_\theta + W w_s^i \mathbf{e}_s. \tag{10}$$

As a consequence, the boundary condition (8c) can be written in the form

$$w_r^i = w_\theta^i = w_s^i = 0 \quad \text{on} \quad r = 1. \tag{11}$$

In this cylindrical coordinate system the continuity equation, (8a), can be written in the form

$$\frac{U}{\delta}\frac{1}{r}\frac{\partial}{\partial r}\left(r w_r^i\right) + \frac{V}{\delta}\frac{1}{r}\frac{\partial w_\theta^i}{\partial \theta} + W\frac{\partial w_s^i}{\partial s} = 0. \tag{12}$$

Here, to enforce conservation of mass, $U = V = \delta W$ is employed. Integrating (12) over the capillary cross-section

$$\begin{aligned} 0 &= W \int_0^{2\pi} \int_0^1 \left[ \frac{1}{r}\frac{\partial}{\partial r}\left(r w_r^i\right) + \frac{1}{r}\frac{\partial w_\theta^i}{\partial \theta} + \frac{1}{r}\frac{\partial w_s^i}{\partial s} \right] r dr d\theta, \\ &= W \int_0^{2\pi} \left[r w_r^i\right]_{r=0}^{r=1} d\theta + W \int_0^1 \left[r w_\theta^i\right]_{\theta=0}^{\theta=2\pi} dr + W \int_0^{2\pi} \int_0^1 \frac{\partial w_s^i}{\partial s} dr d\theta. \end{aligned} \tag{13}$$

Applying the no-slip boundary condition, (8c), and enforcing continuity of $w_\theta^i$ at $\theta = 0$ and $2\pi$, it was deduced that

$$w_s^i = w_s^i(r, \theta). \tag{14}$$

As such, for each component, the momentum equation, (8b), can be written in the form

$$\begin{aligned} \mathbf{e}_i \cdot \mathbf{e}_r &= \frac{1}{\delta}\frac{\partial P^i}{\partial r} \\ &\quad - \mu_{abs}\frac{W}{\delta}\left[\frac{1}{r}\frac{\partial}{\partial r}\left(r\frac{\partial w_r^i}{\partial r}\right) + \frac{1}{r^2}\frac{\partial^2 w_r^i}{\partial \theta^2} - \frac{w_r}{r^2} - \frac{2}{r^2}\frac{\partial w_\theta}{\partial \theta} + \delta^2 \frac{\partial^2 w_r^i}{\partial s^2}\right], \end{aligned} \tag{15a}$$

$$\begin{aligned} \mathbf{e}_i \cdot \mathbf{e}_\theta &= \frac{1}{\delta r}\frac{\partial P^i}{\partial \theta} \\ &\quad - \mu_{abs}\frac{W}{\delta}\left[\frac{1}{r}\frac{\partial}{\partial r}\left(r\frac{\partial w_\theta^i}{\partial r}\right) + \frac{1}{r^2}\frac{\partial^2 w_\theta^i}{\partial \theta^2} - \frac{w_\theta}{r^2} + \frac{2}{r^2}\frac{\partial w_r}{\partial \theta} + \delta^2 \frac{\partial^2 w_\theta^i}{\partial s^2}\right], \end{aligned} \tag{15b}$$

$$\mathbf{e}_i \cdot \mathbf{e}_s = \frac{\partial P^i}{\partial s} - \mu_{abs}\frac{W}{\delta^2}\left[\frac{1}{r}\frac{\partial}{\partial r}\left(r\frac{\partial w_s^i}{\partial r}\right) + \frac{1}{r^2}\frac{\partial^2 w_s^i}{\partial \theta^2}\right]. \tag{15c}$$

Poiseuille's Law specifies that flow along the $s$-axis has a parabolic profile based on the radial coordinate $r$ and the pressure gradient along $s$. Therefore, the scaling of $W = \delta^2$ was chosen to retain this profile to leading-order. Consequently, the final scaling of $\mathbf{w}^i$ is given by

$$\mathbf{w}^i = \delta^3 w_r^i \mathbf{e}_r + \delta^3 w_\theta^i \mathbf{e}_\theta + \delta^2 w_s^i \mathbf{e}_s, \tag{16}$$

and so the system (15) reduces to

$$\begin{aligned}\mathbf{e}_i \cdot \mathbf{e}_r &= \frac{1}{\delta}\frac{\partial P^i}{\partial r} \\ &\quad - \mu_{abs}\delta\left[\frac{1}{r}\frac{\partial}{\partial r}\left(r\frac{\partial w_r^i}{\partial r}\right) + \frac{1}{r^2}\frac{\partial^2 w_r^i}{\partial \theta^2} - \frac{w_r}{r^2} - \frac{2}{r^2}\frac{\partial w_\theta}{\partial \theta} + \delta^2\frac{\partial^2 w_r^i}{\partial s^2}\right],\end{aligned} \tag{17a}$$

$$\begin{aligned}\mathbf{e}_i \cdot \mathbf{e}_\theta &= \frac{1}{\delta r}\frac{\partial P^i}{\partial \theta} \\ &\quad - \mu_{abs}\delta\left[\frac{1}{r}\frac{\partial}{\partial r}\left(r\frac{\partial w_\theta^i}{\partial r}\right) + \frac{1}{r^2}\frac{\partial^2 w_\theta^i}{\partial \theta^2} - \frac{w_\theta}{r^2} + \frac{2}{r^2}\frac{\partial w_r}{\partial \theta} + \delta^2\frac{\partial^2 w_\theta^i}{\partial s^2}\right],\end{aligned} \tag{17b}$$

$$\mathbf{e}_i \cdot \mathbf{e}_s = \frac{\partial P^i}{\partial s} - \mu_{abs}\left[\frac{1}{r}\frac{\partial}{\partial r}\left(r\frac{\partial w_s^i}{\partial r}\right) + \frac{1}{r^2}\frac{\partial^2 w_s^i}{\partial \theta^2}\right]. \tag{17c}$$

Equating coefficients of $\mathcal{O}(\delta^0)$ in (17a) and (17b) gives,

$$0 = \frac{1}{r}\frac{\partial P^i}{\partial r} + \mathcal{O}(\delta) \quad \text{and} \quad 0 = \frac{1}{r}\frac{\partial P^i}{\partial \theta} + \mathcal{O}(\delta), \tag{18}$$

which implies that $P^i = P^i(s)$, to leading-order. Finally, (17c) becomes

$$\mu_{abs}\left[\frac{1}{r}\frac{\partial}{\partial r}\left(r\frac{\partial w_s^i}{\partial r}\right) + \frac{1}{r^2}\frac{\partial^2 w_s^i}{\partial \theta^2}\right] = \frac{dP^i}{ds} - \mathbf{e}_i \cdot \mathbf{e}_s + \mathcal{O}(\delta). \tag{19}$$

As $P^i$ is independent of both $r$ and $\theta$ to leading-order, $w_s^i$ is of the form

$$w_s^i(r,\theta) = \frac{1}{\mu_{abs}}\left(\frac{dP^i}{ds} - \mathbf{e}_i \cdot \mathbf{e}_s\right) f(r,\theta). \tag{20}$$

Noting, that since $w_s^i$ is independent of $s$ and so

$$\frac{dP^i}{ds} - \mathbf{e}_i \cdot \mathbf{e}_s = h, \tag{21}$$

where $h$ is a constant.

The function $f(r,\theta)$ satisfies

$$\frac{1}{r}\frac{\partial}{\partial r}\left(r\frac{\partial f}{\partial r}\right) + \frac{1}{r^2}\frac{\partial^2 f}{\partial \theta^2} = 1 \quad \text{for} \quad 0 \le r \le 1, \tag{22}$$

and subject to the boundary condition

$$f = 0 \quad \text{on} \quad r = 1. \tag{23}$$

Seeking a radially symmetric solution for $f$ and one that is finite at $r = 0$ leads to

$$w_s^i(r) = \frac{\delta^2}{4\mu_{abs}} \left( \frac{dP^i}{ds} - \mathbf{e}_i \cdot \mathbf{e}_s \right) \left( r^2 - 1 \right) \quad \text{for} \quad 0 \le r \le 1. \tag{24}$$

Therefore,

$$\mathbf{w}^i(r) = \frac{\delta^2}{4\mu_{abs}} \left( \frac{dP^i}{ds} - \mathbf{e}_i \cdot \mathbf{e}_s \right) \left( r^2 - 1 \right) \mathbf{e}_s + \mathcal{O}(\delta^3) \quad \text{for} \quad 0 \le r \le 1. \tag{25}$$

Re-dimensionalising and neglecting higher-order terms leads to

$$\mathbf{w}^i(R) = \frac{1}{4\mu_{abs}} \left( \frac{dP^i}{ds} - \mathbf{e}_i \cdot \mathbf{e}_s \right) \left( R^2 - a^2 \right) \mathbf{e}_s + \mathcal{O}(\delta^3) \quad \text{for} \quad 0 \le R \le a, \tag{26}$$

where $R$, $P^i$, $\mu_{abs}$ and $s$ are all dimensional variables and so $\left[\mathbf{w}^i\right] = \mu\text{m}^3\,\text{s/kg}$ where $\left[P^i\right] = \mu m$.

Finally, the micro-cell flux, $\mathbf{q}^i$ (with units $\mu\text{m}^5\,\text{s/kg}$) can be computed by integrating (26) over the capillary cross-section,

$$\begin{aligned} \mathbf{q}^i &= \int_0^{2\pi} \int_0^a \mathbf{w}^i(R)\, R\, dR\, d\theta, \\ &= -\frac{\pi d^4}{128\mu_{abs}} \left( \frac{dP^i}{ds} - \mathbf{e}_i \cdot \mathbf{e}_s \right) \mathbf{e}_s, \end{aligned} \tag{27}$$

where $d = 2a$ is the capillary diameter. Equation (27) represents a modified Poiseuille law, due to an extra forcing term $-\mathbf{e}_j \cdot \mathbf{e}_s$, as a consequence of the the tissue-scale pressure gradient.

### Appendix E: $\beta$-Optimisation

Numerical optimisation is required to conserve mass in the continuum domain. Here, the sum of the arteriolar and venular tree inflows and outflows, via the discrete set of sources and sinks, must be equal to the known net flow into the continuum domain, $q_{act}$. To ensure conservation of mass, a Newton-Raphson procedure was employed to minimise the function $Q = q_{sum} - q_{act}$, through the drainage parameter, $\beta$. Following (34) in the manuscript, the sum of source flow, $q_{sum}$ is given by

$$q_{sum} = \sum_{i \in N_s} q_i^s = \sum_{i \in N_s} \sum_{j \in N_s} \left( M_{ij}^{net} + M_{ij}^{cap} \right)^{-1} \left( p_j^b - p_c \right), \tag{28}$$

which defines the total inflow (negative if outflow) into the capillary domain supplied by the discrete vasculature.

Writing the continuum matrix defined by (31), as function of $\beta$,

$$M_{ij}^{cap} = \frac{1}{\beta}\tilde{M}_{ij}^{cap} \quad \text{for} \quad i,j \in N_s, \tag{29}$$

the minimisation function $Q$ can be written as

$$Q = \sum_{i\in N_s}\sum_{j\in N_s}\left(M_{ij}^{net} + \frac{1}{\beta}\tilde{M}_{ij}^{cap}\right)^{-1}\left(p_j^{base} - p_c\right) - q_{act}. \tag{30}$$

The derivative of $Q$ with respect to $\beta$ yields

$$\frac{dQ}{d\beta} = \frac{1}{\beta^2}\sum_{i\in N_s}\sum_{j\in N_s}\sum_{k\in N_s}\left(M_{ij}^{net} + \frac{1}{\beta}\tilde{M}_{ij}^{cap}\right)^{-1}\cdot \tilde{M}_{jk}^{cap}\cdot q_k^s. \tag{31}$$

Due to the order of $\beta$ we introduce $y = \log_{10}\beta$, and so the minimisation of $Q(\beta)$ is performed by updating the $y$ iteratively according to

$$y_{n+1} = y_n - \frac{1}{\beta\ln(10)}Q(\beta_n)\cdot\left(\frac{dQ(\beta_n)}{d\beta}\right)^{-1}, \tag{32}$$

where $n$ is the current iteration, $Q$ is defined by (30) and $dQ/d\beta$ by (31). Equation (32) is used to update the value of $y$, starting from an initial value of $y = -10^{-3}$ until $Q$ is less than 2.5% of total inflow into the tissue volume or a maximum number of iterations is reached.

### References


[1] Paul W. Sweeney, Angela D'Esposito, Simon Walker-Samuel, and Rebecca J. Shipley. Modelling the transport of fluid through heterogeneous, whole tumours in silico. *PLOS Computational Biology*, 15(6): e1006751, 6 2019. ISSN 1553-7358. doi: 10.1371/journal.pcbi.1006751.

[2] Amy Smith. *Multi-scale modelling of blood flow in the coronary microcirculation.* PhD thesis, University of Oxford, 2013.

[3] Paul W. Sweeney. *Realistic numerical image-based modelling of biological tissue substrates.* PhD thesis, University College London, 2018. URL `http://discovery.ucl.ac.uk/id/eprint/10049410`.

[4] Rebecca J Shipley and S Jonathan Chapman. Multiscale modelling of fluid and drug transport in vascular

tumours. *Bulletin of mathematical biology*, 72(6):1464–91, 8 2010. ISSN 1522-9602. doi: 10.1007/s11538-010-9504-9.

[5] A R Pries and T W Secomb. Microvascular blood viscosity in vivo and the endothelial surface layer. *American journal of physiology. Heart and circulatory physiology*, 289(6):H2657–H2664, 2005. ISSN 0363-6135. doi: 10.1152/ajpheart.00297.2005.

[6] Rebecca J Shipley. *Multiscale modelling of fluid and drug transport in vascular tumours*. PhD thesis, University of Oxford, 2008.
